# Condensin loop extrusion properties, roadblocks, and role in homology search in *S. cerevisiae*

**DOI:** 10.1101/2024.09.12.612585

**Authors:** Vinciane Piveteau, Chloé Dupont, Hossein Salari, Agnès Dumont, Jérôme Savocco, Daniel Jost, Aurèle Piazza

**Affiliations:** Université de Lyon, ENS de Lyon, Université Claude Bernard, CNRS UMR5239, Laboratoire de Biologie et Modélisation de la Cellule, 46 Allée d’Italie, 69007 Lyon, France

**Keywords:** Condensin, loop extrusion, SMC, DNA double-strand break, homology search, mating-type switching

## Abstract

The *in vivo* mechanism, *cis*-acting roadblocks, and biological functions of loop extrusion by eukaryotic SMC complexes are incompletely defined. Here, we identify condensin-dependent Hi-C contact stripes at the Recombination Enhancer (*RE*) and the *rDNA* in *S. cerevisiae*. The *RE* is an autonomous condensin loading site only active in *MAT*a cells from which oriented, unidirectional loop extrusion proceeds with an estimated processivity ∼150-250 kb and a density ∼0.04-0.18 that varies across the cell cycle. Centromeres, replication forks and highly-transcribed RNA PolII-dependent genes are roadblocks for condensin. Cohesin is not an obstacle for condensin while Top2 promotes its loop extrusion activity. A DNA double-strand break at *MAT* blocks loop extrusion, resulting in the establishment of a ∼170 kb-long *RE*-*MAT* loop. The *RE* and the DSB are required and sufficient to form this site-specific loop, which promotes *RE*-proximal homology identification in the early stages of recombinational DNA break repair. We propose that the juxtaposition of the broken *MAT*a site and its target *HML*α donor is the relevant structure by which condensin promotes *MAT*a-to-α switching.

## Introduction

Most Structural Maintenance of Chromosome (SMC) family of ATPase complexes, including cohesin, condensin, and Smc5/6, share the ability to extrude DNA loops *in vitro* presumably via a conserved mechanism (Ganji *et al*, 2018; Davidson *et al*, 2019; Kim *et al*, 2019; Kong *et al*, 2020; Kim *et al*, 2020; Golfier *et al*, 2020; Pradhan *et al*, 2022; Shaltiel *et al*, 2022; Ryu *et al*, 2022; Pradhan *et al*, 2023; Janissen *et al*, 2024). Studying the function and activities of these complexes in cells has proven more challenging, as they generally do not have well-defined loading sites from which extrusion can be experimentally tracked. This is at the notable exception of the condensin-like bacterial SMC loading at *parS* sites (Gruber & Errington, 2009; Sullivan *et al*, 2009), and of a specialized condensin complex (condensin DC) loaded at *rex* sites on the X chromosome in XX hermaphrodites *C. elegans* animals (Morao *et al*, 2022), which enabled studying the loop extrusion activities of these SMCs and identify roadblocks *in vivo* with Hi-C (Morao *et al*, 2022; Brandão *et al*, 2021, 2019; Wang *et al*, 2017; Tran *et al*, 2017).

Condensin I and II are conserved SMC complexes specialized in chromosome assembly and segregation (Hirano, 2016). In *S. cerevisiae*, a single condensin complex individualizes mitotic chromosomes and promotes rDNA segregation at anaphase (Lazar-Stefanita *et al*, 2017; Renshaw *et al*, 2010; Guérin *et al*, 2019; Freeman *et al*, 2000). It is also associated with chromatin at all cell cycle phases, suggesting roles outside of mitosis (Freeman *et al*, 2000; D’Ambrosio *et al*, 2008; Leonard *et al*, 2015). Recently, it has been involved in mating-type (*MAT*) switching (Li *et al*, 2019; Dinda *et al*, 2023), a genetic event induced right prior or upon S-phase entry and that takes ∼1 hour to complete (Hicks *et al*, 1977; Nasmyth, 1983; Connolly *et al*, 1988). Mating-type (*MAT*) switching is a specialized recombination-dependent process that must select between two competing silent donor loci. As such, it represents a classic model for studying the mechanisms underlying the establishment of specific interactions along chromosomes. *MAT* switching is initiated by a site-specific DNA double-strand break (DSB) inflicted by the homothallic (HO) endonuclease at the *MAT* locus on chr. III (Haber, 2012). Gene conversion occurs using one of two silent donors carrying the opposite mating-type information: *HML*α and *HMR*a at the left and right extremity of chr. III, respectively. Of note, *MAT* is located ∼95 kb away from *HMR*a on the same chromosome arm, yet *MAT*a cells efficiently use the more distant *HML*α donor present ∼186 kb away across the centromere. Donor preference is regulated by a bipartite “recombination enhancer” (*RE*) element present ∼14 kb away from *HML*α, only active in *MAT*a cells (Wu & Haber, 1996). The first part binds the Rad51-ssDNA filament in *trans* and promote homology search in its vicinity (Renkawitz *et al*, 2013; Dumont *et al*, 2024), and the second part is required for the horseshoe folding of chr. III, that brings the left and right chromosomal arms in close proximity (Dekker *et al*, 2002; Belton *et al*, 2015; Li *et al*, 2019). This horseshoe folding depends on condensin, which is enriched at the *RE* in a Sir2-, Tof2-, Fob1-, cohibin-, and Mcm1-dependent manner only in *MAT*a cells, suggesting that it is loaded there (Li *et al*, 2019; Dinda *et al*, 2023). Accordingly, condensin and the proteins required for its enrichment at the *RE* promote usage of *HML*α upon DSB formation at *MAT*a (Li *et al*, 2019; Dinda *et al*, 2023). Such non-mitotic role of condensin has been ascribed to its chr. III folding activity. A second site enriched for condensin is the replication fork barrier (*RFB*) site within *rDNA* repeats, with largely overlapping requirements with the *RE* at the notable exception of the Matα2-repressible Mcm1 factor (Johzuka & Horiuchi, 2009). These putative loading sites represent an opportunity to study the loop extrusion activity of condensin in eukaryotic cells and its function in establishing specific chromosomal interactions.

Here, using Hi-C, we show the existence of discrete condensin-dependent contact stripes emanating from the *RE* and the *rDNA* locus, and provide several lines of evidence that condensin unidirectionally extrudes chromatin loops from these sites with a defined orientation in cells. In conjunction with a polymer model for loop extrusion, we exploit this system to infer the basic loop extrusion properties of condensin *in vivo* and delineate its regulation in *cis* by various types of roadblocks. Functionally, condensin loop extrusion properties are uniquely exploited during *MAT*a-to-α switching, upon DSB-dependent reconfiguration of chr. III structure.

## Results

### Two condensin-dependent contact stripes in the budding yeast genome

Chromatin immunoprecipitation (ChIP)-based assays revealed that the *RE* is the unique genomic region most enriched for condensin in *MAT*a cells (**Fig. 1A**) (Li *et al*, 2019; Costantino *et al*, 2020; Rossi *et al*, 2021). Such approaches cannot straightforwardly disambiguate loading and accumulation of long-range translocases such as SMCs, unlike Hi-C (Wang *et al*, 2017; Brandão *et al*, 2021; Wike *et al*, 2021; Guo *et al*, 2022; Morao *et al*, 2022; Kim *et al*, 2025). Previous Hi-C experiments failed to clearly established such contact signature at the *RE* (Belton *et al*, 2015; Li *et al*, 2019) while recent micro-C data indicated the presence of a contact stripe (Dinda *et al*, 2023). Our Hi-C protocol that yields contact maps at sub-kilobase resolution was applied on an asynchronous population of haploid *MAT*a and *MAT*α cells. It confirmed the presence of a discrete, *MAT*a-specific contact stripe emanating from the *RE* and stretching across chr. III (**Fig. 1A**). It depended on the presence of the *RE*, as did the overall chr. III folding (**Fig. EV1A**) (Belton *et al*, 2015). This stripe could be visualized as a 4C-like profile and compared to control sites in the yeast genome (**Fig. 1B** and see **Methods** for the selection of control sites), which revealed a specific enrichment of contacts from 40 kb to the end of chr. III. These observations suggested that a heterogeneous population of loops link the *RE* and the rest of chr. III specifically in *MAT*a cells (Belton *et al*, 2015; Li *et al*, 2019; Dinda *et al*, 2023; Li *et al*, 2024). Similar to the *RE*, a contact stripe could be observed emanating from the left rDNA-flanking region and extending up to the centromere of chr. XII, exhibiting similar contact enrichment features independently of the mating type (**Fig. EV1B**).

**Figure 1:**
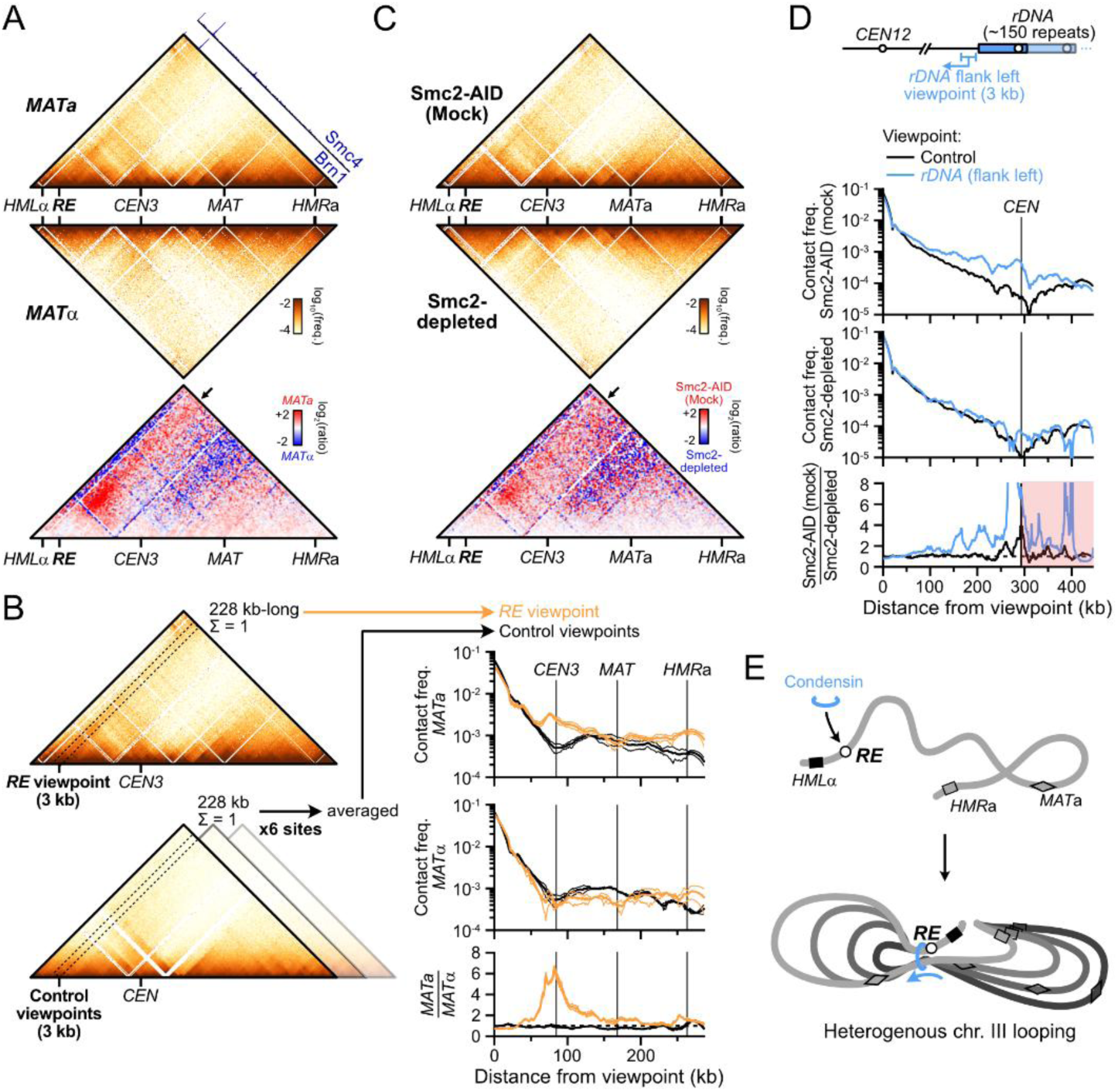
Two condensin-dependent Hi-C contact stripes in the budding yeast genome. (A) Hi-C contact maps (top) and ratio map (bottom) of chr. III in exponential cultures of *MAT*a (APY142) and *MAT*α (APY295) cells. Data show the merge of n=4 and n=2 biological replicates, respectively. Calibrated ChIP-Exo profiles are from ref. (Rossi *et al*, 2021). Bin: 1 kb. (B) Left: Rationale for the computation of *RE* 4C-like profiles and comparison to equivalent control sites (for precise genomic coordinates, see **Methods**). Right: 4C-like contact profiles of the *RE* and of the average of 6 control sites in *MAT*a and *MAT*α cells, from Hi-C data in (A). Bottom: Ratio of *MAT*a and *MAT*α profiles. Data show mean ± SEM of n=4 and 2 biological replicates, respectively. (C) Hi-C contact maps and ratio maps of Smc2-AID and Smc2-depleted *MAT*a cells (YTG155). Bin: 1 kb. (D) 4C-like profiles with the left rDNA-flanking region as a viewpoint, and the average of 6 corresponding control sites. Data show n=1 biological replicate. (E) Model for *RE*-dependent chr. III loop folding by condensin in *MAT*a cells.

We addressed the dependency of these stripes on condensin by depleting its Smc2 subunit using the auxin-inducible degron (AID) system (Morawska & Ulrich, 2013) in an asynchronous culture of haploid *MAT*a cells (**Fig. 1C** and **EV1C**). Condensin depletion did not affect the cell cycle distribution, the genome-wide probability of contact P_c_ as a function of genomic distance *s*, and only exhibited substantial changes in contacts on chr. III and XII (**Fig. EV1C-E**) (Lazar-Stefanita *et al*, 2017; Costantino *et al*, 2020). The associated stripe on chr. III was lost upon condensin depletion (**Fig. 1C** and **EV1F**). The contact stripe emanating from the rDNA-flanking region and extending up to the centromere of chr. XII also depended on condensin (**Fig. 1D** and **EV1E**). No corresponding stripe was found on the telomere side of the rDNA (**Fig. EV1E**), suggesting that loop formation by condensin is oriented. No other condensin-dependent stripe was detected genome-wide.

These results show that condensin forms two loops with well-defined anchors on chr. III and XII. In particular, condensin folds chr. III has an heterogenous population of loops from the *RE* specifically in *MAT*a cells (**Fig. 1E**), corroborating prior suggestions (Li *et al*, 2019; Dinda *et al*, 2023). *MAT*α cells thus provide a physiological “off” state for condensin specifically at the *RE*, which we exploited in the following sections.

### The *RE* is a directional and oriented condensin loop anchor

The *RE* contains two main modules: a left Fkh1-binding region and a right condensin-binding region (**Fig. 2A**)) (Li *et al*, 2019; Haber, 2012). In order (i) to address whether the *RE* was sufficient to form a contact stripe outside of the context of chr. III, (ii) to determine its reliance on the Fkh1 module, and (iii) to disambiguate the stripe signal from that of the brush configuration conferred by the Rabl chromosome organization, we introduced the full *RE* or a truncated version lacking the Fkh1 module (*RE-right*) at an interstitial location in the longest chromosome arm of the yeast genome (right arm of chr. IV, **Fig. 2A-B** and **EV2A**). It resulted in the formation of a single, *MAT*a-specific stripe emanating from the *RE* or *RE-right* constructs that extended up to the right end of chr. IV, several hundreds of kilobases away (**Fig. 2B-C** and **EV2A**). The direction of the stripe could be reversed upon inversion of the *RE* constructs (**Fig. 2B, D** and **EV2A**). Hence, the *RE* is required and sufficient for forming a discrete contact stripe exhibiting a specific orientation in *MAT*a cells, independently of its Fkh1-binding module.

**Figure 2:**
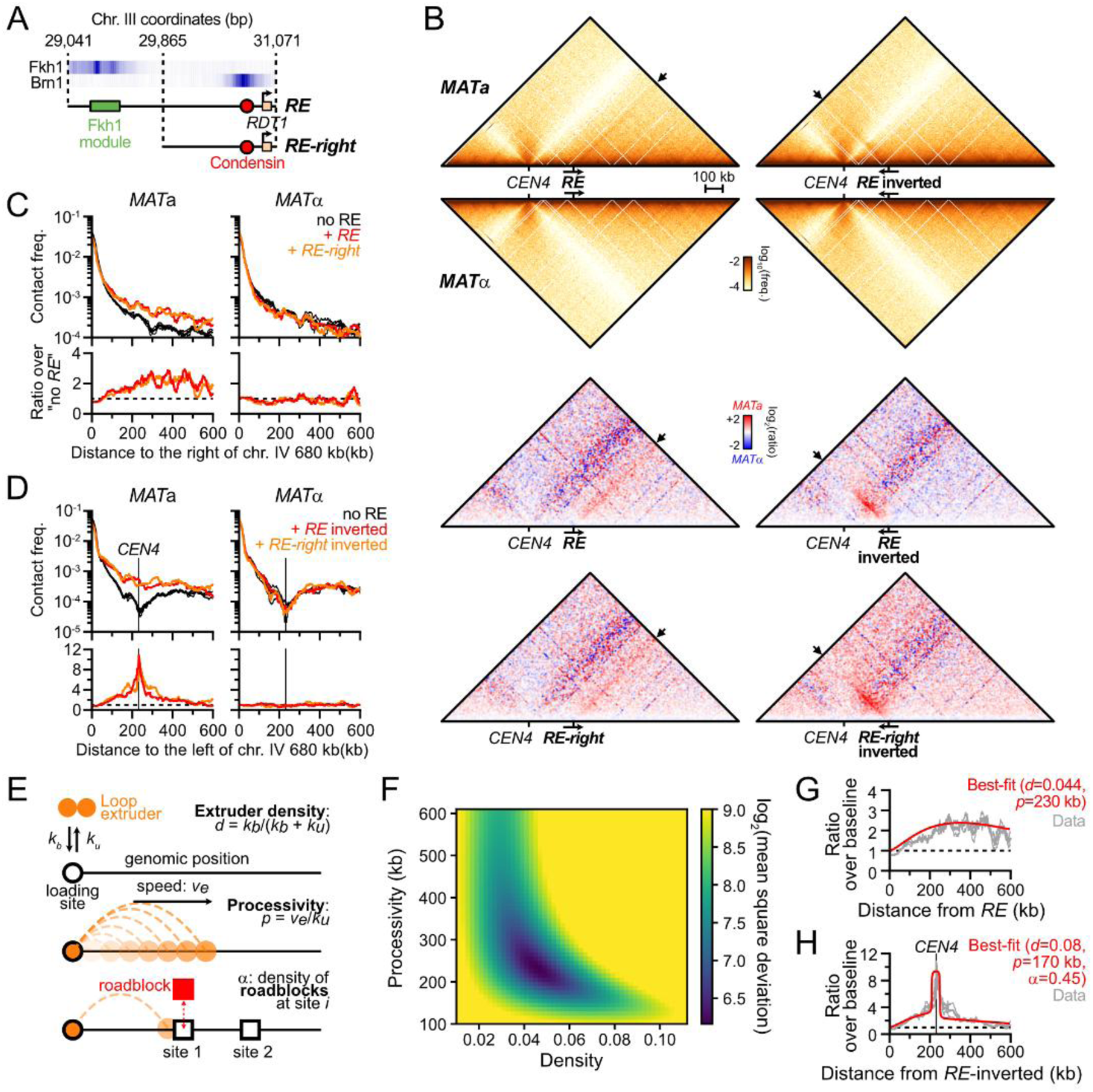
Condensin loop extrusion properties. (A) Description of the *RE* and *RE-right* constructs introduced at position 680,258 on chr. IV. ChIP profiles are from (Rossi *et al*, 2021). (B) The *RE* is autonomous in forming a single oriented contact stripe. Top: Hi-C contact maps of chr. IV in *MAT*a and *MAT*α cells bearing the *RE* at position 680,258 either in the forward (APY1846 and APY1848) or inverted (APY1911 and APY1913) orientation. Bottom: Ratio maps of *MAT*a over *MAT*α cells in cells bearing either the *RE* or *RE-right* in the forward or the inverted orientation. The *RE-right* data are from **Fig. EV2A**. Hi-C maps are binned at 5 kb. Ratio maps are binned at 10 kb. (C) Top: 4C-like contact profiles of the chr. IV 680 kb coordinate either unmodified, or upon insertion of the *RE* or the *RE-right* constructs in *MAT*a (APY266, APY1846 and APY1850) and *MAT*α cells (APY295, APY1848 and APY1852). Bottom: Ratio of the 4C-like profiles with a *RE* construct over the “no *RE*” baseline. Data show the mean ± SEM of n=4 and 2 biological replicates for the “no *RE*” *MAT*a and *MAT*α cells, respectively. n=1 biological replicate for each +*RE* constructs. (D) Same as (C) with *RE* constructs in the inverted orientation (APY1848, APY1852, APY2058, and APY2060), and 4C profiles oriented towards the left end-side of chr. IV. (E) Rationale of the unidirectional loop extrusion crunching model, parameters, and output. (F) Heat map of the mean squared deviation (MSD) between theoretical predictions and experimental data, expressed as the ratio of Hi-C interactions of the *RE* and *RE-right* constructs inserted on chr. IV in the forward direction versus no *RE* in *MAT*a cells (see **Fig. EV2B**). The region with the lowest MSD corresponds to the best fit of processivity and loop density. (G) Observed data and best-fit simulated contact frequencies. Data are the average of the *RE* and *RE-right* constructs in the forward orientation in *MAT*a cells from (C). (H) Same as (C) with *RE*-inverted and *RE-right*-inverted construct in *MAT*a cells (data from (D)).

These observations provide strong evidence that condensin extrudes chromatin loops in *S. cerevisiae* cells. These loops have two well-defined anchors that correspond to highly enriched condensin binding sites: the *RE* and at the *rDNA*. They may originate from an unidirectional loop extrusion process initiated at the site present at the base of the stripe; or from a site-specific block of loop extrusion events initiated at multiple dispersed sites (Fudenberg *et al*, 2016, 2017; Vian *et al*, 2018). In the following sections we assumed the first scenario to be correct. This assumption is supported by additional evidence presented below and summarized in the **Discussion**.

### Density and processivity of loop extrusion by condensin inferred from biophysical modelling

We sought to infer properties of loop extrusion by condensin in cells by comparing experimental contact data with the output of a quantitative “scrunching” framework (**Fig. 2E**). It integrates the unidimensional stochastic motion of loop extruders along the chromosome into a simple polymer model to predict the impact of loop extrusion on contact frequencies, which can be compared to Hi-C contact data (Alipour & Marko, 2012; Sanborn *et al*, 2015; Fudenberg *et al*, 2016; Chan & Rubinstein, 2023). In this framework, condensin can bind and unbind from its loading site at rates *k*_*b*_ and *k*_*u*_, respectively. Upon binding, one leg of the condensin remains anchored at the binding site while the other translocates along the chromosome at a speed *v*_*e*_. These parameters define two key observables: the processivity *p* = *v*_*e*_/*k*_*u*_, which describes the average loop size extruded by condensin in the absence of roadblock; and the density of loop extruders bound to DNA at any single time *d* = *k*_*b*_/(*k*_*b*_ + *k*_*u*_). Pausing at site-specific roadblocks *i* can also be modeled. This allows one to estimate the probability of observing an extruding loop between the condensin loading site and any other site along the chromosome and thus to predict their contact frequency. We first applied our framework to infer model parameters consistent with the stripe emanating from the *RE* and *RE-right* constructs inserted on chr. IV that lacks discernable loop extrusion roadblocks (see below). This led to best-fitting parameters for condensin processivity *p* in the ∼170-250 kb range, with a density of condensin on chromatin *d* in the ∼0.04-0.06 range (**Fig. 2F-G** and **EV2B**). Consistently, loop extrusion proceeded with a processivity *p*= 170 kb and a density *d* = 0.08 from the inverted *RE* constructs, and revealed a pause frequency ∼0.45 at the centromere (**Fig. 2H**).

Condensin-mediated loop extrusion proceeded similarly from its endogenous loading sites, with a processivity *p* = 120 kb and a density *d* = 0.11 from the *rDNA* and *p* = 150 kb and a density *d* = 0.06 from the *RE* in asynchronous cells. Consistently, the *SMC2-AID* control strain yielded similar *p* = 150-170 kb and *d* = 0.05-0.07, while Smc2 depletion caused a reduction of *d* = 0.02 at both sites without affecting the processivity. These results indicate that condensin is a chromatin loop extruder with a processivity ∼3- to 9-fold greater than that estimated for cohesin in mitotic and meiotic *S. cerevisiae* cells (Schalbetter *et al*, 2017, 2019).

### The density of condensin-dependent loops is cell cycle-regulated

Although budding yeast condensin is detected on chromatin at all cell-cycle stages, its enrichment at specific genomic sites and its role in chromatin compaction and *rDNA*-*CEN12* tethering varies along the cell cycle (Lazar-Stefanita *et al*, 2017; Freeman *et al*, 2000; D’Ambrosio *et al*, 2008; Leonard *et al*, 2015; Bhalla *et al*, 2002). Consequently, we analyzed the loop extrusion by condensin on chr. III and XII in G1, S-phase, and in cells arrested in G2/M either artificially (*i.e.* repression of *CDC20* or addition of nocodazole) or upon activation of the DNA damage checkpoint (*i.e.* formation of an unrepairable site-specific DNA double strand break on chr. V), and determined best-fitting loop extrusion parameters for each condition (**Fig. 3A-E** and **EV2A-F**). The contact stripes emanating from the *RE* on chr. III and the rDNA on chr. XII were detectable at all cell cycle stages, with phase-specific variations in intensity and pausing at centromeres (**Fig. 3A-E** and **EV2B-F**, and see below).

**Figure 3:**
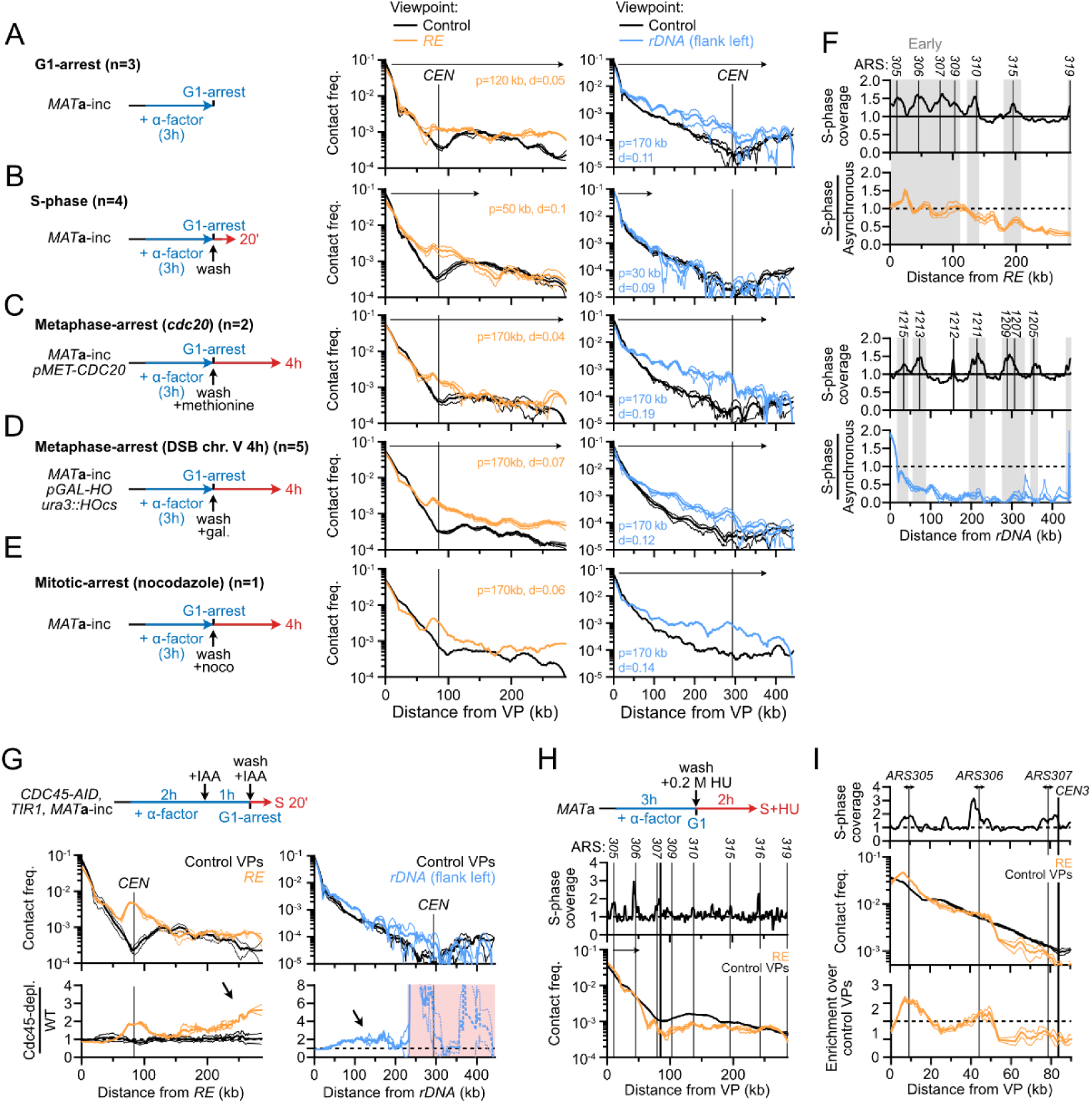
Loop extrusion by condensin is inhibited by replication forks. (A-E) *RE* and *rDNA*-flanking 4C-like contact profiles (A) upon G1-arrest (APY266), (B) during S-phase (APY539 and APY607, merged), (C) upon metaphase-arrest due to *CDC20* repression (APY537), (D) upon DDC-induced metaphase-arrest due to formation of a single unrepairable HO-induced DSB at *ura3* (APY266), and (E) upon mitotic-arrest in the presence of nocodazole (APY266). All cells are *MAT*a. Best-fitting simulation processivity (*p*) and density (*d*) values are indicated. Data show mean ± SEM. The number of biological replicates n is indicated on each panel. (F) Top: Coverage from Hi-C reads in S-phase. Bottom: Ratio of the contact stripes in S-phase over the average of asynchronous cultures at the *RE* (top) and left rDNA-flanking (bottom) viewpoints. From data in (B). (G) Entry in S-phase in the absence of replication origin firing (Cdc45-AID + IAA) partially restores loop extrusion by condensin on chr. III and XII (APY539). Data show mean ± SEM of n=2 biological replicates from (D’Asaro *et al*, 2025). (H) Stopping replisome progression immediately after S-phase entry prevents loop extrusion by condensin. Data show mean ± SEM of n=3 biological replicates from (Jeppsson *et al*, 2022). (I) Zoom on data in (C).

Cells arrested in G1 with alpha-factor exhibited a stripe signal similar to that of asynchronous cells on both chr. III and chr. XII (**Fig. 3A** and **EV2B**). Consistently, best-fit parameters yielded *p* = 170 kb and *d* = 0.11 for the *rDNA* and *p* = 120 kb and *d* = 0.05 for the *RE*. Differently, the contact stripe on chr. III during S-phase failed to progress past ∼120 kb (**Fig. 3B** and **EV2C**), resulting in an altered chr. III conformation in which contacts between the left arm and the right arm distal to *MAT* strongly decreased (**Fig. EV2G**). Likewise, the condensin extrusion span on chr. XII did not exceed ∼80 kb.

Simulations indicated a reduction of processivity (*p* = 80 kb and 30 kb at the *RE* and the *rDNA*, respectively) while loop density remained unchanged (*d* = 0.09-0.1). Hence, condensin translocation, but not loading, is inhibited during S-phase. Finally, the contact stripes were overall similar in G2/M and asynchronous cells on both chr. III and XII (**Fig. 3C-E** and **S2D-F**): the estimated processivity remained constant (*p* = 170 kb) while loop density *d* increased modestly at the *rDNA*, between 0.12 to 0.18 from 0.11 in G1. Hence, loop extrusion by condensin at these specific sites is active in mitosis prior to anaphase and presumably regulated at the loading stage, while its overall loop extrusion activity remains unaltered (Lazar-Stefanita *et al*, 2017; Guérin *et al*, 2019; Freeman *et al*, 2000; Leonard *et al*, 2015; Strunnikov *et al*, 1995).

### Replication forks stall loop extrusion by condensin

The condensin loop extrusion span was strongly reduced on both chr. III and XII during S-phase (**Fig. 3B**). *RE*-bound loop extrusion initiates in an early-replicated region and progressed on chr. III up to the broad vicinity of *ARS310*, which corresponds to the first late replicating region encountered (**Fig. 3F**). Differently, the condensin loop extrusion collapsed before encountering the first early-replicating *ARS1215* region immediately on chr. XII. In order to address whether replication forks are roadblocks for condensin, we depleted the DNA replication initiation factor Cdc45 in G1-arrested cells and released them in S-phase (**Fig. 3G**). Preventing origins from firing partially restored condensin progression up to the end of chr. III and over ∼ 200 kb on chr. XII, although not to levels observed in G1 and mitosis (**Fig. 3G**). Oppositely, authorizing replication firing but immediately blocking replication forks progression at high hydroxyurea (HU) concentration further reduced the condensin loop extrusion span (**Fig. 3H** and **EV2H-I**). The immediate blocking of replication forks upon HU treatment further allowed visualizing discrete loops between the *RE* and replication origins (**Fig. 3I**). These observations indicate that replication forks are strong condensin translocation roadblocks, independently of replisome progression. The incomplete restoration of condensin loop extrusion upon Cdc45 depletion further suggests that DDK-primed replisome components can also act as roadblock for condensin, independently of the establishment of replication forks.

### Condensin-dependent loops are shortened in the absence of Top2

Top2 promotes loop extrusion by Condensin DC in *C. elegans* (Morao *et al*, 2022) and condensin function in chromosome compaction is tightly coupled to Top2 activity in bacteria and eukaryotes (Hirano, 2016). We thus addressed whether condensin translocation requires Top2 activity by re-analyzing published Hi-C datasets of cells arrested in mitosis upon treatment with benomyl in which Top2 was either conditionally inactivated at restrictive temperature (*top2-4* (Holm *et al*, 1985)) or depleted (Top2-AID) upon auxin addition prior to S-phase entry and up to mitotic arrest (Jeppsson *et al*, 2024) (**Fig. EV4A-D**). The span of loop extrusion by condensin was reduced to ∼80 kb on chr. III and seemingly abolished on chr. XII in the absence of Top2 activity (**Fig. EV4A-D**). The phenotype was slightly more penetrant upon Top2 heat inactivation than Top2 depletion. These results suggest that Top2 promotes loop extrusion by condensin, and that the extent of this effect is locus-specific.

### Determinants of the condensin roadblock at centromeres

To gain insights into the nature and determinants of the condensin loop extrusion pausing at centromeres (*e.g.* **Fig. 1A** and **2B, H**), we related the roadblock at *CEN3* to the folding of the peri-centromeric regions across the cell cycle (**Fig. 4A**). No clear roadblock was detected in G1, consistent with the lack of condensin enrichment at peri-centromeres at that cell cycle stage (Leonard *et al*, 2015). A narrow roadblock appeared in S-phase, and became highly prominent in metaphase-arrested cells. This roadblock at the centromere was still observed upon treatment of cells with the microtubule-depolymerizing drug nocodazole, indicating it is independent of the mitotic spindle and of centromere clustering (**Fig. 4B**). Furthermore, Cdc45-depleted cells that reach metaphase in the absence of replication, and thus lack the sister chromatid required for the spindle to exert tension, also retained a localized roadblock (**Fig. 4C**). Roadblock appearance coincided with S-phase, when loop-extruding cohesins are loaded (Costantino *et al*, 2020; Dauban *et al*, 2020) and restructure the peri-centromere (Yeh *et al*, 2008; Paldi *et al*, 2020). The role of cohesin in blocking condensin was addressed by re-analyzing S-phase and metaphase Hi-C data obtained after Scc1 depletion in G1-arrested *MAT*a cells (**Fig. EV5A-B**) (Dumont *et al*, 2024; D’Asaro *et al*, 2025). The condensin roadblock persisted at the centromere (**Fig. 4D** and **EV5C**). More generally, condensin-dependent stripes appeared unaffected upon Scc1 depletion, indicating that condensin is oblivious to the presence of cohesin on chromatin (**Fig. EV5A-B**). Consequently, the centromeric roadblock for condensin is independent of spindle tension, centromere clustering, the centromeric chromatin hairpin, cohesin, and centromere replication.

**Figure 4:**
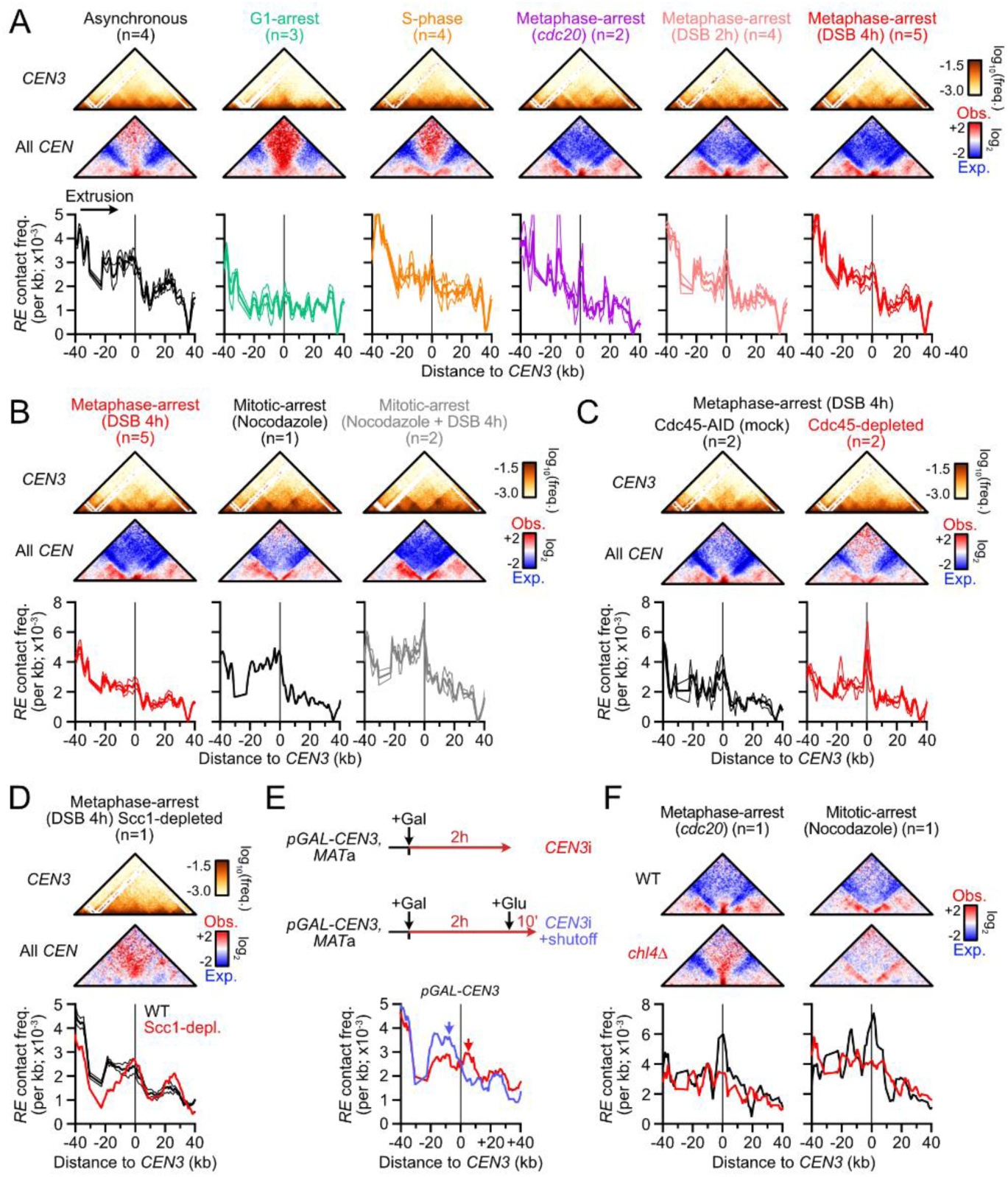
Centromeres stall condensin translocation in a kinetochore-dependent manner. (A) Condensin roadblock at *CEN3* across the cell cycle. Top: Hi-C maps of the *CEN3*-surrounding region. Middle: Aggregated contact maps of all centromeres. Bin: 1 kb. Bottom: 4C-like profile centered on the *CEN3* region with the *RE* as a viewpoint. Data show mean ± SEM of zooms of **Fig. 2A-E**. n: number of biological replicates. Data show mean ± SEM. (B) 4C-like profile with the *RE* as a viewpoint in cells arrested at mitosis in the presence or absence of nocodazole. n: number of biological replicates. Data show mean ± SEM. (C) 4C-like profile with the *RE* as a viewpoint in metaphase-arrested cells with replicated chromatids (Cdc45-AID mock) or non-replicated chromatids (Cdc45-depleted prior to S-phase entry; APY513). Data show mean ± SEM of n=2 biological replicates from (Dumont *et al*, 2024). (D) 4C-like profile with the *RE* as a viewpoint in metaphase-arrested with or without cohesin (Scc1-depleted, APY1481). Data show n=1 biological replicate from (Dumont *et al*, 2024). (E) Top: *CEN3* inactivation scheme (APY1745). Bottom: 4C-like profile with the *RE* as a viewpoint in the *CEN3*-surrounding region. n=1 biological replicate. (F) Aggregated contact maps of all centromeres (top) and 4C-like profile with the *RE* as a viewpoint (bottom) in WT and *chl4Δ* cells arrested in metaphase or in mitosis. Data show n=1 biological replicate each from (Paldi *et al*, 2020).

We then addressed whether the centromeric function was required to block condensin in *cis*. To this end, we specifically inactivated the centromere of chr. III upon inducible transcription across *CEN3* using the *pGAL-CEN* system (**Fig. 4E**) (Hill & Bloom, 1987). Such system causes the loss of the centromere-specific Cse4-containing nucleosome, and thus of the kinetochore, from *CEN3* (Nakabayashi & Seki, 2022). *CEN2* inactivation was used as a control (**Fig. EV5D-E**). Strong transcriptional activation for 2 hours upon galactose addition caused a specific ∼4-fold loss of contacts between the transcribed centromere and the 15 other centromeres, which was not restored after a 10 minutes transcriptional shutoff with glucose (**Fig. EV5D**). The appearance and disappearance of the border pattern typical of highly-transcribed genes (Banigan *et al*, 2023; Salari *et al*, 2024; Chapard *et al*, 2025) confirmed the efficient transcriptional activation and shutoff (**Fig. EV5E**). The centromeric roadblock was abolished upon *CEN3* transcription and rapidly recovered following transcriptional shutoff (**Fig. 4E**). Transcription across *CEN2* did not affect the roadblock at *CEN3* (**Fig. EV5F**), indicating that the effect observed upon *CEN3* inactivation occurs in *cis*. Furthermore, the roadblock was lost in a strain defective for the dispensable outer kinetochore component Chl4, both in the presence and absence of spindle tension (**Fig. 4F**). These results indicate that the condensin roadblock at the centromere depends on the presence of the kinetochore in *cis*, irrespective of its role in mediating spindle tension and centromere clustering.

### Highly-transcribed RNA PolII genes transiently stall condensin translocation

Besides centromeres, the contact stripes on chr. III and XII exhibited local contact variations suggestive of the presence of additional roadblocks along chromosome arms, which could be highlighted upon detrending of the contact stripes (**Fig. EV6A**). These positions were compared to various genomic and chromatin features detected by ChIP-Exo conducted in rich, glucose-containing media (Rossi *et al*, 2021) (**Fig. 5A**). Clearly, loop extrusion paused immediately ahead of sites enriched for RNA PolII, which exhibit the conserved “boundary” Hi-C pattern typical of highly-transcribed genes (**Fig. 5A**) (Banigan *et al*, 2023; Salari *et al*, 2024; Chapard *et al*, 2025). These sites were also enriched for condensin, consistent with it pausing at highly-transcribed genes (**Fig. 5A**). These pause profiles were also observed in G1-and metaphase-arrested cells (**Fig. EV6B**). The pause site ∼160 kb away from the rDNA and not associated with a RNA PolII-enriched site (**Fig. 5A**) corresponded to the region immediately upstream of the *GAL2* gene, specifically activated in our galactose-containing culture conditions (**Fig. 5A-B**; galactose did not change the expression of the other genes in the *CEN12-rDNA* interval (see (Pelechano *et al*, 2013) and **Fig. EV6D** below). Accordingly, the boundary Hi-C pattern is readily detected at *GAL2* and other *GAL* genes (**Fig. 5A-B** and **EV6C**). Cells grown in the absence of galactose did not exhibit the boundary nor the increased *rDNA-GAL2* looping (**Fig. 5B-C** and **EV6D**). Transcriptional induction by addition of galactose for only 10 minutes was sufficient to induce transcription of the *GAL* genes (Brouwer *et al*, 2023) and enrich for the *rDNA-GAL2* loop (**Fig. 5C** and **EV6D-E**). Conversely, repression of *GAL* genes upon glucose addition to galactose-containing media for 10 minutes (Johnston *et al*, 1994) caused a partial loss of the *rDNA-GAL2* loop (**Fig. 5D** and **EV6F**). Hence, condensin pausing ahead of a highly-transcribed PolII gene can be rapidly established and reversed, suggesting that it is a primary consequence of its transcriptional activity. Pausing occurred irrespectively of the gene orientation relative to the directionality of loop extrusion (*e.g. PDC1* and *PGK1* vs. *AHP1* and *GAL2* in **Fig. 5A**). Inactivation of RNA PolII, either upon heat-inactivation of its Rpb1 subunit or upon thiolutin treatment, exerted profound effects on chromosome organization (Jeppsson *et al*, 2022), which prevented us from determining the role of transcription in regulating the processivity of loop extrusion by condensin over large chromosomal segments.

**Figure 5:**
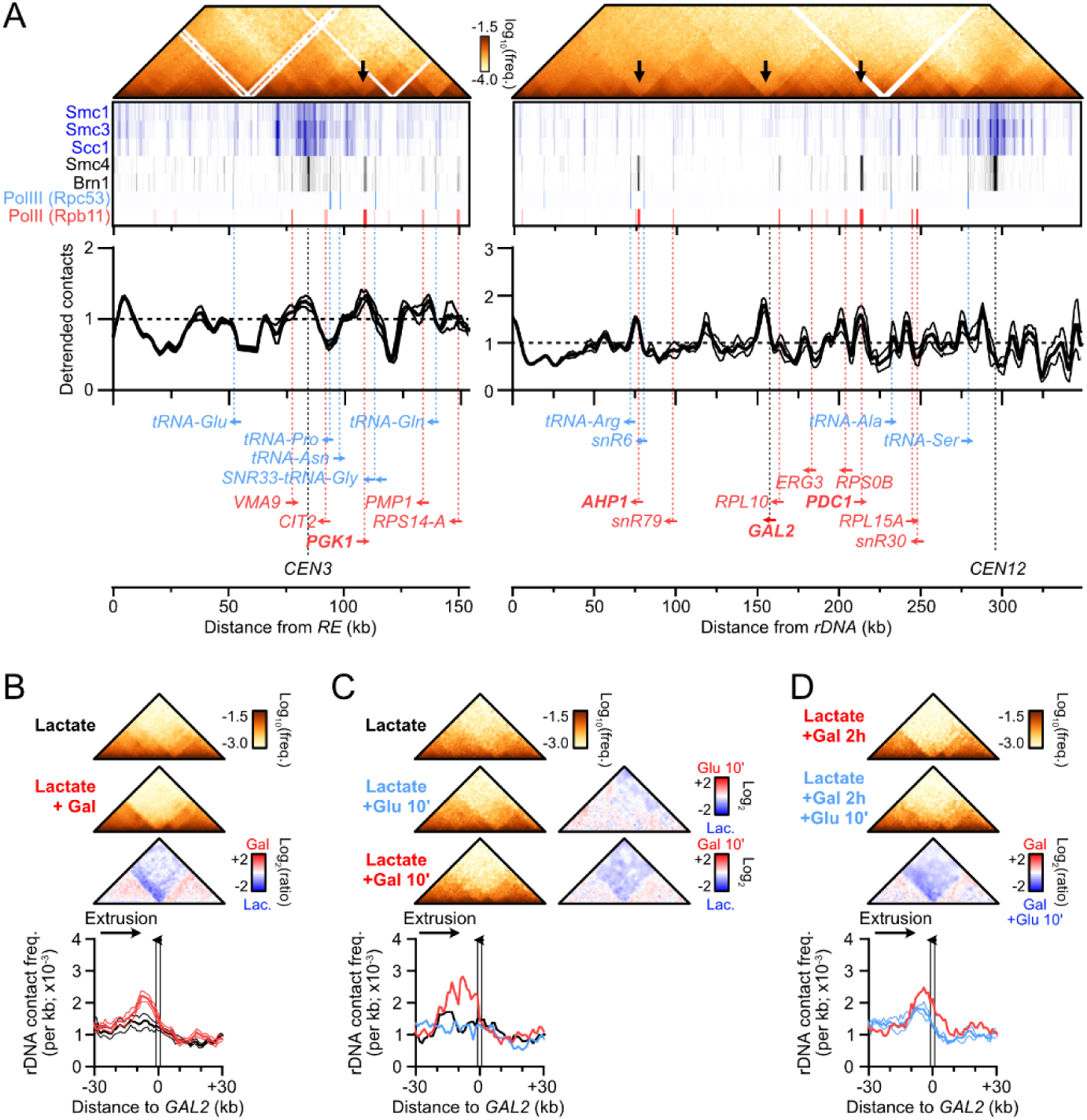
Highly transcribed RNA PolII-dependent genes stall condensin translocation. (A) From top to bottom, Hi-C contact maps, ChIP-Exo enrichment profiles of cohesin, condensin, RNA PolIII, and RNA PolII subunits (from (Rossi *et al*, 2021)), detrended *RE* and rDNA-flanking 4C-like profiles in asynchronous *MAT*a cells (n=4), and relevant genomic features. Contact data were obtained from cells grown in galactose-containing media while ChIP-Exo data were obtained in glucose-containing media, explaining the discrepancy at *GAL2*. (B) Hi-C matrices (top) and 4C-like profiles with the left rDNA-flanking region as a viewpoint (bottom) in the 60 kb region surrounding the *GAL2* gene in asynchronous *MAT*a cells (APY266) grown in the presence of lactate or lactate supplemented with galactose. Data show mean ± SEM of n=5 and 6 biological replicates, respectively. Bin: 1 kb. (C) Same as (B) in *MAT*a cells (APY142) grown in lactate media and supplemented or not with 2% galactose or 2% glucose for 10 minutes. Data show n=1 biological replicate. (D) Same as (B) with samples used in Fig. 4E.

We further noted the presence of a pause site ∼120 kb from the rDNA locus (position ∼330 kb) not associated with a high occupancy of RNA PolII or PolIII (**Fig. 5A** and **EV6B**). The nature of this roadblock remains to be determined.

### Rad51-independent reconfiguration of chr. III structure upon DSB at *MAT*

What is the functional significance of the condensin-mediated loop extrusion spanning across chr. III? Previous reports revealed that condensin promoted usage of the *RE*-proximal *HML*α donor during *MAT* switching in *MAT*a cells, which was presumed to originate from its role in chr. III folding (Li *et al*, 2019; Dinda *et al*, 2023). Differently, we sought to address whether a DSB induced at *MAT* may act as a condensin roadblock, with the resulting *RE-MAT* loop directly juxtaposing the DSB to the *HML*α donor. To this end, we induced *HO* over-expression from a galactose-inducible promoter in asynchronous *MAT*a and *MAT*α cells. Repair-deficient *rad51Δ* mutants were used to (i) ensure the maintenance of a homogeneous population of *MAT*a or *MAT*α cells (*i.e.* no switching), and (ii) prevent the Rad51-dependent preferential recruitment of the DSB to the *RE* in *trans* (Renkawitz *et al*, 2013; Dumont *et al*, 2024). The efficiency of DSB induction, inferred from Hi-C coverage, was similar in all strains assayed (**Fig. EV7A**).

DSB induction resulted in the formation of a ∼170 kb-long *RE*-*MAT* loop only in *MAT*a cells (**Fig. 6A-C**). Specifically, the contact stripe emanating from the *RE* was interrupted at the level of the DSB in the ∼20 kb region immediately upstream of the DSB while contacts downstream were not different from control sites, indicating an absence of loop extrusion past the DSB (**Fig. 6A-C**, and see below). A *MAT*a strain with an unrepairable DSB induced at *ura3* on chr. V did not exhibit such arrest, showing that the DSB exerts its roadblock effect in *cis*. Deletion of the Fkh1-containing left part of the *RE* did not affect formation of the DSB-induced *RE*-*MAT*a loop (**Fig. 6D, E** and **EV7B, C**), indicating that the condensin-containing right part of the *RE* is sufficient for loop formation. Inversion of this *RE-right* element abolished loop formation (**Fig. 6D-E**), establishing that the directional translocation of condensin towards the DSB is required to form the *RE*-*MAT*a loop. These observations show that a DSB at *MAT*a blocks the condensin-mediated loop extrusion initiated at the *RE* in *cis*, converting a heterogeneous population of loops along chr. III into a site-specific *RE*-DSB loop (**Fig. 6F**).

**Figure 6:**
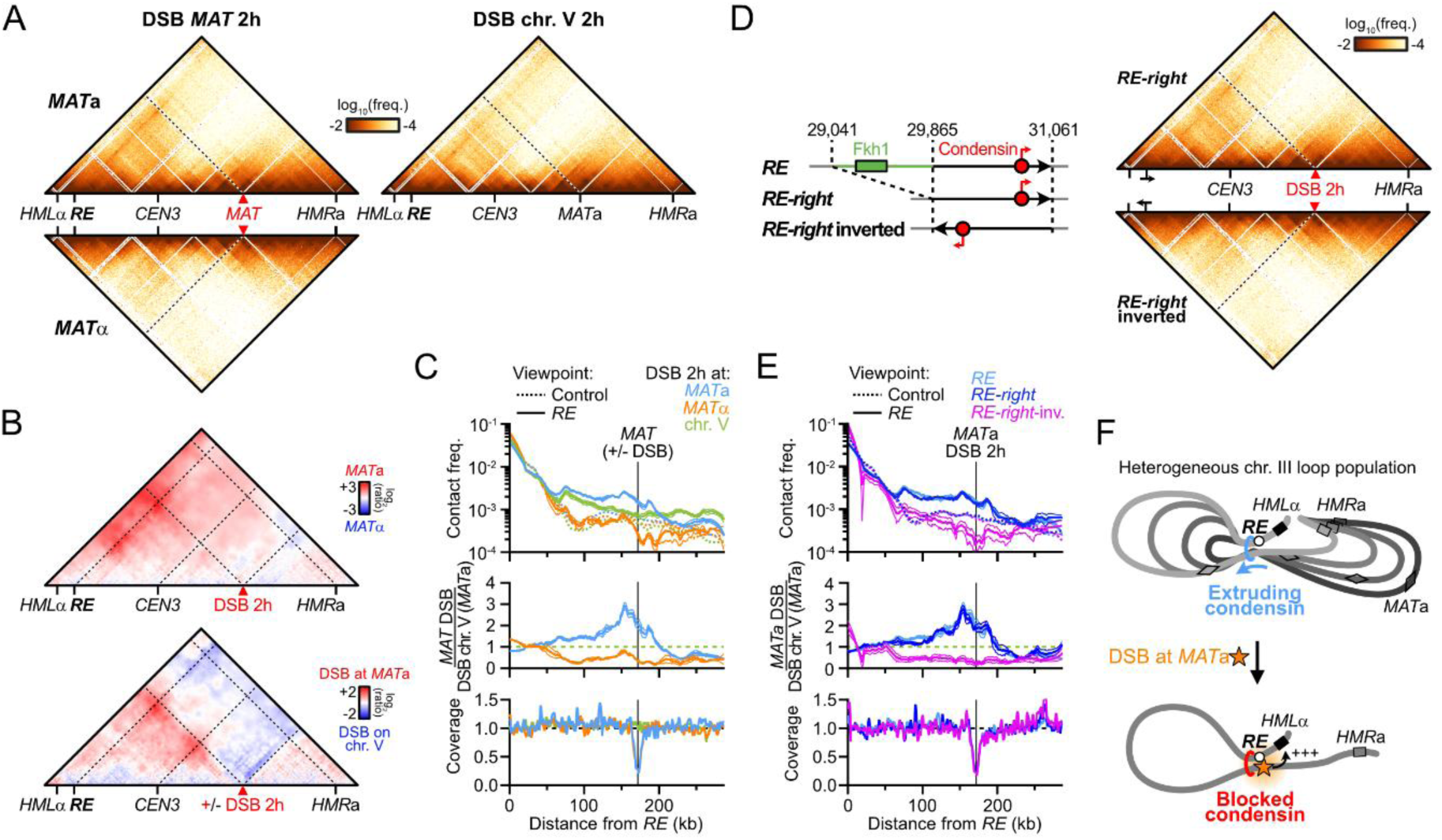
A Rad51-independent *RE*-*MAT*a loop forms upon DSB induction. (A) Hi-C contact maps in *MAT*a and *MAT*α *rad51*Δ cells (APY1267 and APY1264, respectively) at 2 hours post-DSB induction at *MAT* (n=4 and 2 biological replicates, respectively). A *MAT*a strain with an unrepairable DSB on chr. V (APY266) is shown for comparison 2 hours post-DSB induction (n=4 biological replicates). Bin: 1 kb. (B) Ratio maps of data in (A). (C) Top: 4C-like profiles with the *RE* (or 6 control sites) as a viewpoint, from data in (A). Middle: Ratio of the 4C-profiles over that of a *MAT*a cell with a DSB on chr. V. Bottom: Coverage from Hi-C reads showing the resection tract length at *MAT* (see also **Fig. EV7A**). Data show mean ± SEM. (D) Left: Scheme of the *RE-right* and *RE-right*-inverted constructs. Right: Hi-C contact maps of *MAT*a *rad51*Δ cells with the *RE-right* and *RE-right*-inverted constructs (APY2072 and APY2079). Data show merged of n=3 biological replicates each. Bin: 1 kb. (E) Top: 4C-like profiles with the *RE*, *RE-right* and *RE-right*-inverted constructs (or 6 control sites) as a viewpoint, from data in (D). Middle: Ratio of the 4C-profiles over that of a *MAT*a cell with a DSB on chr. V. Bottom: Coverage from Hi-C reads showing the resection tract length at *MAT* (see also **Fig. EV7B**). Data show mean ± SEM. (F) Model for chr. III structure reconfiguration by condensin upon DSB formation into a homogeneous *RE-MAT*a loop. This structure juxtaposes the DSB and the *HML*α donor.

The *RE*-*MAT* loop resulted in broken *MAT*a engaging *HML*α ∼3.2-fold more than *HMR*a (**Fig. EV7D-F**). It is opposite to the situation in *MAT*α cells where *HMR*a is preferred ∼2-fold over *HML*α, leading to a ∼6.4-fold difference for donor preference between both mating-types upon DSB formation (**Fig. EV7F**). This donor preference is only ∼1.7-fold in the absence of a DSB (**Fig. EV7D, F**). Consequently, the structure that drives preferential interaction between *MAT*a and its target *HML*α donor is not the heterogeneous loop folding observed in asynchronous *MAT*a cells, but the *RE*-*MAT* loop specifically formed upon DSB formation. This site-specific loop is likely the relevant structure in promoting *MAT*a-to-α switching (see below, **Fig. 6F** and **Discussion**) (Li *et al*, 2019; Dinda *et al*, 2023).

Intriguingly, a stripe emanating from the *RE* and stretching up to the DSB site appeared 4 hours post-DSB induction in *MAT*α cells, bringing the DSB in close proximity to *HML*α (**Fig. EV7G-H**). This loop only formed after the elimination of the *MAT*α genes by resection (**Fig. EV7A**), whose α2 gene product represses the Mcm1-mediated condensin loading at the *RE* (Li *et al*, 2019; Dinda *et al*, 2023). Hence, the active repression of condensin loading at the *RE* in *MAT*α cells may provide a back-up redirection of homology search towards *HML*α in case of a prolonged failure to repair a *MAT*α DSB using the primary *HMR*a donor (**Fig. EV7I**).

### The *RE* and a DSB are sufficient to establish a directional loop

In order to address whether other elements on chr. III participate in *cis* in the establishment of the *RE*-DSB loop and to ascertain its formation in a Rad51-proficient background, we induced a single unrepairable DSB 165 kb downstream of the *RE-right* construct on chr. IV in *MAT*a and *MAT*α WT cells (**Fig. 7A, B** and **EV8A**; the endogenous HO cut-site at *MAT* was inactivated by a single point mutation). This minimal system recapitulated the loop observed on chr. III (**Fig. 7A-C**). Inverting the *RE-right* construct abolished loop formation (**Fig. 7A, B**), demonstrating that the *RE*-DSB loop is formed as a consequence of condensin translocating towards the DSB in *cis*. Hence, formation of a *RE*-DSB loop reflects a basic property of condensin encountering a DSB that can operate outside of the natural context of chr. III and *MAT*, and in a Rad51-proficient context. The pause frequency of condensin at the DSB was estimated to be ∼90% in this simple system (**Fig. 7C**), confirming the strong or absolute roadblock a DSB represents for condensin in cells.

**Figure 7:**
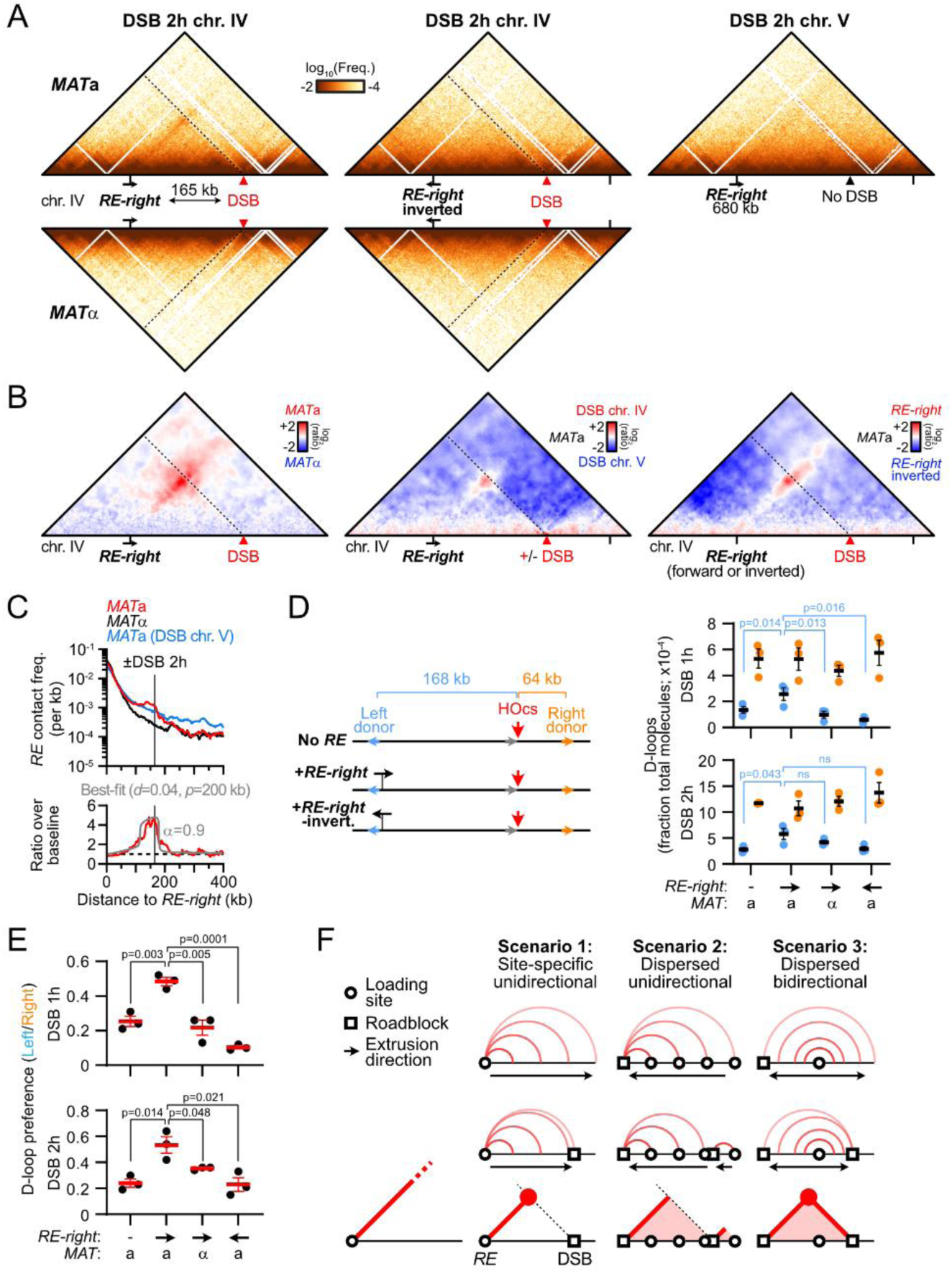
The *RE*-DSB loop is portable and promotes *RE-*proximal homology search. (A) Hi-C contact maps of the chr. IV region spanning from 555 to 970 kb that contains the *RE*-*right* constructs at position 680 kb and an unrepairable DSB at position 845 kb in *MAT*a (APY1850 and APY2058) and *MAT*α cells (APY1852 and APY2060), as well as in a *MAT*a strain with an unrepairable DSB on chr. V (APY1918). Data show n=1 biological replicate. Bin: 2 kb. (B) Ratio maps of data in (D). (C) Top: 4C-like profiles from *RE-right* on chr. IV as a viewpoint, from data in (A). Bottom: Observed data and best-fit simulated contact frequencies. Data represent the ratio of *MAT*a over *MAT*α data. (D) Left: donor competition system to address the role of the *RE*-DSB loop in biasing homology search. Right: Absolute D-loop levels at the left and right donors at 1 and 2 hours post-DSB induction in strains without (APY2083) or with the *RE-right* (*MAT*a: APY2085, *MAT*α: APY2087) or *RE-right*-inverted (APY2088) inserted near the left donor. Data show individual biological replicates (n=3) and mean ± SEM. P-values were obtained using a paired ratio Student *t* test. None of the comparisons for the right donor are significant. (E) Donor preference, computed from data in (D). Data show individual biological replicates (n=3) and mean ± SEM. P-values were obtained using a Student *t* test. (F) Predicted contact patterns with different loop extrusion scenarios in presence of a DSB roadblock. The observed data in (B) correspond to scenario 1.

### The *RE-*DSB loop promotes *RE*-proximal homology identification

To address the functional relevance of this condensin-dependent *RE*-DSB loop during recombinational DNA break repair, we quantified D-loops formed by the left DSB end at two competing intra-chromosomal donors positioned on each side of the DSB in our minimal experimental system on chr. IV (**Fig. 7D**). The donors were introduced at intergenic locations and at positions and distances from the DSB emulating that of *HML*α and *HMR*a relative to *MAT* on chr. III (∼168 kb to the left and ∼64 kb to the right of the DSB site introduced at position 845,464, respectively). The *RE-right* construct was further introduced near the left donor in the forward or inverted orientation, only the former of which leads to the formation of a *RE*-DSB loop (**Fig. 7A-B**). Absolute D-loop levels at each donor were quantified 1 and 2 hours post-DSB induction using the D-loop-Capture assay (Piazza *et al*, 2019; Reitz *et al*, 2022; Djeghmoum & Piazza, 2025) (**Fig. 7D** and **EV8B-C**). D-loop formation was biased 4-fold towards the most proximal right donor (**Fig. 7D-E**). Introduction of the *RE-right* near the left donor halved this preference by promoting D-loop formation at that donor, while having no effect on D-loops formed at the right donor (**Fig. 7D-E** and **EV8D**). No such stimulation was observed if condensin extruded loops away from the DSB (*RE-right*-inverted) or if loop extrusion from the *RE-right* site was prevented in *MAT*α cells (**Fig. 7E**). Hence, homology identification at the *RE*-proximal donor is stimulated only in the context in which a condensin-dependent *RE*-DSB loop can be formed. Condensin whose translocation has been blocked at a DSB thus promotes homology search around its loading site.

## Discussion

Here, we provided evidence that condensin extrudes chromatin loops from two specific sites in the budding yeast genome and determined several of its loop extrusion properties and roadblocks in cells. Notably, condensin forms a long-range loop structure between its loading site and a DSB, which promotes homology search at the loop anchor. We propose a revised model of donor selection during mating-type switching that exploits the specificities of loop extrusion by condensin we defined here.

### Properties of loop extrusion by condensin in S. cerevisiae

Contact stripes detected by Hi-C correspond to a heterogeneous population of loops sharing a discrete anchor. Such loops may be formed by a unidirectional extrusion process initiated at a loading site present at the base of the stripe (scenario 1), or by a site-specific block for unidirectional or bi-directional loop extrusion processes initiated at dispersed sites within the genomic interval covered by the stripe (scenario 2 and 3, respectively; **Fig. 7F**) (Fudenberg *et al*, 2016, 2017; Vian *et al*, 2018). Here, we present evidence in favor of the first “loading” scenario. Indeed, in the “blocking” scenarios, the numerous on-going loop extrusion events that have not yet reached the blocking site are expected to nonetheless link distant sites along the main Hi-C diagonal. Condensin removal should thus affect the probability of contact P_c_ as a function of genomic distance *s*, which was not observed (**Fig. EV1D**). This absence of an effect on the genome-wide P_c_(*s*) could not be explained either by distributed loading events only on chr. III and the *CEN12-rDNA* intervals, as stripes could also be observed by introducing the *RE* on chr. IV (**Fig. 2B**). Importantly, the *RE-*DSB loops were only associated with a single stripe emanating from the *RE,* and not from the DSB (**Fig. 6A-B** and **7A-B**). This observation is incompatible with scenario 2 and 3, which both predict the appearance of stripes anchored at the DSB. These multiple lines of evidence indicate that condensin is loaded at the *RE*, from which it unidirectionally extrudes loops, as proposed in scenario 1 (**Fig. 7D**). This model is consistent with the unidirectional loop extrusion activity of condensin on naked DNA reported *in vitro* (Ganji *et al*, 2018; Kim *et al*, 2020; Shaltiel *et al*, 2022; Analikwu *et al*, 2025).

Intriguingly, loop extrusion by condensin from its loading sites has a defined orientation, leading to a single stripe, as observed upon introduction of the *RE* on chr. IV (**Fig. 2B-D** and **EV2A, B**). The mechanism specifying this orientation remains to be determined.

Condensin extrudes loops with a processivity ∼150-250 kb that does not substantially vary between the G1 and G2/M phases of the cell cycle, but is reduced during S-phase. Condensin-mediated loop density is modestly decreased in G1, consistent with its overall reduced presence on chromatin and the lower amount of its limiting Ycg1 component at that cell cycle phase (D’Ambrosio *et al*, 2008; Leonard *et al*, 2015; Doughty *et al*, 2016). Finally, Top2 activity is required for condensin-mediated loop extrusion (**Fig. EV4**), similar to the X-specific condensin DC in *C. elegans* (Morao *et al*, 2022). It suggests a broad conservation of the coupling between the strand passage activity of Top2 and the loop extrusion activity of condensin, which likely underlies their shared function in chromatid condensation and decatenation during mitosis (Wood & Earnshaw, 1990; Hirano, 2016; Goloborodko *et al*, 2016; Racko *et al*, 2018; Orlandini *et al*, 2019; Dyson *et al*, 2021).

### Roadblocs for loop extrusion by condensin

Arrays of the high-affinity DNA-binding protein Rap1, naturally present at budding yeast telomeres, have been reported to block loop extrusion by condensin *in vitro* and in cells (Guérin *et al*, 2019; Analikwu *et al*, 2025). Here we identified several additional natural roadblocks of varying intensity for condensin-mediated loop extrusion and delineated their main requirements:

- Centromeres are permeable roadblocks in a kinetochore-dependent, but microtubule-, cohesin-, sister chromatid- and thus tension-independent manner.
- RNA PolII genes are weak roadblocks, detected only at the most highly-transcribed genes in both the G1 and G2/M phases of the cell cycle. These roadblocks can be induced and shut off within 10 minutes in *S. cerevisiae*, suggesting that RNA PolII activity itself, or its immediate consequences on chromatin composition and structure, hinders loop extrusion by condensin. The conservation of this roadblock in fission yeast (Lebreton *et al*, 2024) and *B. subtilis* (Gruber & Errington, 2009; Brandão *et al*, 2019) further hints at a basic interference between RNA polymerases and condensin activity, rather than species-specific management of highly-transcribed regions (*e.g.* delocalization to the nuclear periphery).
- Replication forks are strong roadblocks for condensin, independently from their progression. Partial recovery of loop extrusion in S-phase upon depletion of the origin-firing factor Cdc45 suggests that the fork structure itself is an obstacle for condensin translocation. This partial rescue also raises the possibility that specific replisome components loaded at pre-replication complexes could be obstacles outside of the context of forks, as proposed for mouse cohesin and MCM complexes (Dequeker *et al*, 2022). The inability of condensin to overcome replication forks and/or inactive replisome components may contribute to the condensation defects or unreplicated chromosomal regions in mammalian cells (Ono *et al*, 2013; Boteva *et al*, 2020).
- DNA double-strand breaks are strong or absolute condensin roadblocks, independently of Rad51.

Whether these various impediments to condensin loop extrusion involve species-specific protein-protein interactions or whether they are steric in nature, as previously shown with heterologous high-affinity DNA bound elements in *S. cerevisiae* and *in vitro* (Pradhan *et al*, 2022; Analikwu *et al*, 2025), remains to be determined.

### The relevant role of condensin in *MAT*a-to-α switching is to tether the DSB near *HML*α

Condensin loading at the *RE* has previously been shown to promote directional *MAT*a-to-α switching (Li *et al*, 2019; Dinda *et al*, 2023). This modest stimulatory effect was presumed to arise as a consequence of the heterogeneous “horseshoe” folding of chr. III, believed to reduce the average distance between the DSB and its target *HML*α donor in the context of a diffusive 3D search (Coïc *et al*, 2006a; Belton *et al*, 2015; Lassadi *et al*, 2015; Li *et al*, 2019; Dinda *et al*, 2023). Here, we reveal that condensin juxtaposes *MAT*a and the *RE-*surrounding region that includes the *HML*α donor specifically upon DSB formation (**Fig. 6F**), and that such loop promotes homology identification near the *RE* in our minimal *RE*-DSB system on chr. IV (**Fig. 7D-E**). We thus propose that the DSB-induced *RE*-*MAT*a loop is the relevant structure promoting a-to-α switching. In this model, the heterogenous loop population observed in the absence of break represents a futile structure whose role is to poise chr. III for rapid establishment of the *RE*-*MAT* loop upon DSB formation.

Increased proximity between *MAT* and the *RE* has been previously observed cytologically within less than 1 hour of DSB induction, but whether it resulted from the recombination process itself, or whether it preceded it, was not established (Simon *et al*, 2002; Bressan *et al*, 2004; Houston & Broach, 2006; Avşaroğlu *et al*, 2016). Here we show that this association takes place in the absence of Rad51 (**Fig. 6A-C**), indicating that it is not a consequence of homology search or of the Rad51 filament interaction with the Fkh1-containing part of the *RE* (Renkawitz *et al*, 2013; Dumont *et al*, 2024). The condensin-mediated clamping of the DSB and the *RE* is expected to limit their diffusion. Consistently, the diffusion coefficient of the chromatin flanking the *MAT* locus rapidly decreases following DSB induction (Saad *et al*, 2014), unlike DSBs formed on other chromosomes (Dion *et al*, 2012; Miné-Hattab & Rothstein, 2012). Functionally, such clamping should promote the over-sampling of the *RE*-surrounding region and considerably accelerate *HML*α identification. Accordingly, artificial tethering between *HML*α- and *MAT*α-adjacent sites could partially outcompete usage of *HMR*a (Simon *et al*, 2002; Kostriken & Wedeen, 2001), with a magnitude similar to that contributed by condensin in *HML*α usage in *MAT*a cells (Li *et al*, 2019; Dinda *et al*, 2023).

*MAT* switching thus provides a model to study the basic mechanism that establishes selective interactions between chromosomal segments. Biasing for *HML*α usage in *MAT*a cells depends on two main modules in the *RE*: a condensin-loading module whose deletion reduces donor bias by 2-fold (Li *et al*, 2019); and a Fkh1-binding module active only in G2/M whose deletion almost abolishes *HML*α usage (Wu & Haber, 1996; Sun *et al*, 2002; Coïc *et al*, 2006b) (**Fig. 2A**). This major role of Fkh1 at the *RE* specifically depends on its FHA domain (Li *et al*, 2012) and can act in *trans* (Coïc *et al*, 2006b; Lee *et al*, 2016) by recruiting the Rad51-ssDNA filament (Dumont *et al*, 2024; Renkawitz *et al*, 2013). We thus propose that the more modest role of condensin in promoting *MAT*a-to-α switching may be accelerate the establishment of the first link between the *RE* and the broken *MAT*a locus, whose maintenance would subsequently be handed over to Fkh1 and the Rad51-ssDNA filament. This two-step scenario for establishing specific long-range interactions in *cis* along chromosomes thus consists of an initial phase of moderate specificity modulated by intrinsic properties of a SMC complex and its roadblock followed by a maintenance phase dependent on the protein-protein and protein-DNA affinities of two (or more) cognate DNA-bound factors.

### Cohesin and condensin loop extrusion properties differently regulate homology search

All four structurally related SMC complexes (*i.e.* cohesin, condensin, Smc5/6, and Mre11-Rad50-Xrs2) have been implicated in specific or general aspects of recombinational DNA repair in budding yeast. Notably, cohesin regulates homology search during the repair of accidental DNA break in both budding yeast and human cells (Covo *et al*, 2010; Piazza *et al*, 2021; Dumont *et al*, 2024; Teloni *et al*, 2025; Marin-Gonzalez *et al*, 2025) while condensin promotes directional *MAT*a-to-α switching (Li *et al*, 2019; Dinda *et al*, 2023) and, as we show here, homology identification near its loading site (**Fig. 7D-E**).

Mechanistically, cohesin in *S. cerevisiae* and RecN in *C. crescentus* (Piazza *et al*, 2021; Chimthanawala *et al*, 2022; Dumont *et al*, 2024) were proposed to endow the RecA/Rad51-ssDNA filament with the ability to access distant chromatin regions upon directional motion on chromatin in *cis*. To achieve such directional motion, SMCs must anchor at or near the RecA/Rad51-ssDNA filament and thread chromatin unidirectionally from that anchor. In this framework, SMC’s processivity and roadblocks determines the scanning range. Accordingly, we previously showed that cohesin could promote the identification of a donor in *cis* as a function of its processivity, which could be expanded in a *pds5* mutant (Piazza *et al*, 2021). Consistently, the span of RAD51 chromatin enrichment around site-specific DSB sites can be modulated in opposite ways upon depletion of NIPBL^Scc2^ and WAPL (Teloni *et al*, 2025), and the identification of distant ectopic donors in *cis* was reduced in the absence of NIPBL^Scc2^ in human cells (Marin-Gonzalez *et al*, 2025), suggesting a broad conservation of this layer of HR regulation imparted by cohesin.

Differently, condensin promotes a specific interaction between the region surrounding its loading site (*i.e.* the *RE*) and the break site located in its processivity range (**Fig. 6F**). Such conditional *RE*-DSB clamping is made uniquely possible by the properties of loop extrusion by condensin we defined here: its site-specific and oriented loading at the *RE*, its unidirectionality, and its inability to bypass a DSB. Hence, distinct SMC loop extrusion properties are exploited to promote different search strategies in *cis*: long-range scanning by cohesin, and focal search for condensin.

## Methods

### Haploid *S. cerevisiae* strains

Genotypes of *Saccharomyces cerevisiae* (W303 *RAD5*+ background) strains are listed in **Table EV1**. The inducible *HO* expression construct *trp1::pGAL1-HO::hphMX*, the mutagenesis of the endogenous HO cut-site at *MAT* and the DSB-inducible HOcs construct *LY-HOcs* at *ura3* on chr. V and at position 845,464 on chr. IV have been described previously (Piazza *et al*, 2019, 2018, 2021). The *rad51::KanMX* mutation has been obtained by transformation of a PCR fragment amplified from the relevant deletant of the Euroscarf gene deletion collection. The construct *his3::pADH1-OsTIR1-9Myc::HIS3* for OsTir1 E3-ubiquitin ligase expression, as well as the Scc1-V5-AID and Cdc45-FlagX5-AID constructs have been described previously (Piazza *et al*, 2019; Dauban *et al*, 2020). The *SMC2-AID-9Myc::NAT* and *TIR1-9Myc::URA3* constructs were described previously (Guérin *et al*, 2019). The *pGAL1-CEN2* and *pGAL1-CEN3* constructs have been described previously (Hill & Bloom, 1987; Reid *et al*, 2008). The extended recombination enhancer (*RE*) on chr. III (coordinates 29,041-31,071) or its condensin peak-containing right segment (coordinates 29,865-31,071) have been introduced at an inter-genic position on chr. IV (coordinate 680,258) upon CRISPR/Cas9-mediating knock-in using guide 5’-TTGTTTCTACTAATGTGCTG −3’ and repair gene fragments bearing ∼150 bp of homology to each side of the break as described in (Agier *et al*, 2021). The *RE* deletion on chr. III has been obtained by CRISPR/Cas9 as described in (Dumont *et al*, 2024). The deletion of the Fkh1-binding *RE*-*left* region (coordinates 29,041-29,865) has been generated upon CRISPR/Cas9-mediated targeting using two guides (5’-TCTCAAAACCAAATTGCGCA −3’ and 5’-CCAATTCCAAATTCTAGGGA −3’) and a repair fragment bearing ∼150 bp of homology to both sides of the desired deletion junction. The inversion of the condensin-binding *RE-right* region (coordinates 29,866-31,061) has been generated upon CRISPR/Cas9-mediated using two guides (5’-TCTCAAAACCAAATTGCGCA −3’ and 5’-TTGGCTCTATAAAGGAGTTC −3’) and two repair fragments bearing ∼150 bp of homology to both sides of the desired inversion junctions.

Two 556 bp-long donors (corresponding to the position +23 to +578 of the *LYS2* gene) were introduced at position 680,088 and 911,499 on chr. III upon CRISPR/Cas9-mediated targeting using two guides (5’- CAAGATACAAGCCGTTTCCA −3’ and 5’- CGCAATGATGCAATAGTCCA −3’) and two repair fragments containing the donor sequence bearing ∼150 bp of homology to both sides of the desired insertion points. The donor at position 911,499 is flanked by a *TRP1* marker. They bear homology to the left end side of the DSB region introduced at position 845,464 on the same chromosome.

All the coordinates correspond to the S288c R64-2-1 *S. cerevisiae* genome assembly. All the genetic constructs generated in this study are available as annotated Genbank files in **Dataset EV1**.

### Culture media and growth conditions

#### G1-arrest

Exponentially growing cultures in YEP-lactate-galactose (1% yeast extract, 1% peptone, 2% lactate, 2% galactose) were synchronized in G1 by addition of 1 µg/ml alpha-factor (GeneCust) every 30 minutes for 3 hours at 30°C prior to crosslinking.

#### S-phase

Cells arrested in G1 with alpha-factor in YPD medium (1% yeast extract, 1% peptone, 2% glucose) at 30°C were washed 3 times with 50 mL of YPD and released in S-phase at 25°C for 20 minutes prior to crosslinking. For Scc1-AID depletion and Cdc45-AID depletion, 2 mM IAA was added 1 hour prior to, and in all following wash and culture media (described and controlled for by Western blot and FACS in (D’Asaro *et al*, 2025)).

#### Metaphase-arrest (*CDC20* repression)

Metaphase-arrest in strains in which the *CDC20* gene is placed under the control of the *pMET3* promoter (APY537) was performed as in (Dauban *et al*, 2020) with minor media differences. Exponentially growing cells in a supplemented synthetic complete lactate media deprived of methionine (0.67% yeast nitrogen base without amino acids, 2% lactate, and supplemented with a mix of amino-acids lacking methionine) were arrested in G1 upon addition of 1 µg/ml alpha-factor (GeneCust) every 30 minutes for 4 hours at 30°C. Cells were were washed 3 times with 50 mL of YEP-lactate supplemented with 2 mM methionine, and maintained arrested in this media at 30°C for 4 hours prior to crosslinking.

#### Metaphase-arrest (DNA damage checkpoint-induced)

Exponentially growing cultures in YEP-lactate medium of strains bearing the galactose-inducible *pGAL1-HO* construct for expression of the HO endonuclease and its unrepairable *HOcs* target site at *ura3* (APY266) were synchronized in G1 by addition of 1 µg/ml alpha-factor every 30 minutes for 3 hours at 30°C. Cells were washed 3 times with 50 mL of pre-warmed YEP-lactate and released in S-phase in YEP-lactate supplemented with 2% galactose to induce HO expression from the *pGAL1-HO* construct, which targets an unrepairable HOcs at *ura3* on chr. V (Piazza *et al*, 2018, 2019). In one instance, 10 μg/mL nocodazole was added to cause microtubule depolymerization. Cdc45-AID depletion was induced upon addition of 2 mM IAA 1 hour prior to release in S-phase and maintained in all media thereafter. Scc1-AID depletion was induced upon addition of 2 mM IAA upon release in S-phase and HO induction (described and controlled for by Western blot and FACS in (Dumont *et al*, 2024)). Cells were crosslinked at 2 and 4 hours post-DSB induction.

#### Mitotic-arrest (DNA damage checkpoint-induced)

Exponentially growing cultures in YEP-lactate medium at 30°C were supplemented with 10 μg/mL nocodazole and cultured an additional 4 hours prior to crosslinking.

#### Induction of a DNA double-strand break at *MAT*

Exponentially growing Rad51-deficient cells (APY1264 and APY1267) in YEP-lactate medium at 30°C were supplemented with 2% galactose to induce the expression of the HO endonuclease from the *pGAL1-HO* construct. Cells were crosslinked at 2 and 4 hours post-DSB induction.

### Flow cytometry

Approximately 10^7^ cells were collected by centrifugation, re-suspended in 70% ethanol and fixed at 4°C for at least 24h. Cells were pelleted, resuspended in 1 mL of 50 mM sodium citrate pH 7.0 and sonicated 10 seconds on a Bioruptor. After washing, cells were treated with 200 µg of RNase A (Euromedex, cat.9707-C) at 37°C overnight. Cells were then washed and incubated for 30 minutes with 1 mL of 50 mM sodium citrate pH 7.0 with 16 µg of propidium iodide (Fisher Scientific, 11425392). Flow cytometry profiles were obtained on a MACSQuant machine and analyzed using Flowing Software 2.5.1.

### Protein extraction and western blotting

Protein extracts for western blot were prepared from 5.10^7^ to 10^8^ cells. Cells were lysed in cold NaOH buffer (1.85 N NaOH, 7.5% v/v beta-mercaptoethanol) for 10 minutes in ice. Proteins were precipitated upon addition of trichloroacetic acid (15% final) for 10 minutes in ice. After centrifugation at 15,000 g for 5 min, the pellets were resuspended in 100 µL of SB++ buffer (180 mM Tris-HCl pH 6.8, 6.7 M urea, 4.2% SDS, 80 µM EDTA, 1.5% v/v beta-mercaptoethanol, 12.5 µM bromophenol blue). Proteins were denatured upon heating 5 minutes at 65°C. Pre-cleared extracts were resolved on 12% precast polyacrylamide gel (Bio-Rad, cat. 4561043) and blotted on a PVDF membrane (GE Healthcare, cat. 10600023). Membranes were probed with mouse anti-Myc monoclonal antibody (clone 9E11, Invitrogen, cat. MA116637) diluted at 1:1000 for Smc2-Myc-AID, and a mouse anti-GAPDH monoclonal antibody (clone GA1R, Invitrogen, MA515738) diluted at 1:5000. Primary antibodies were revealed with an HRP-conjugated rabbit anti-mouse IgG antibody diluted at 1:5000 (Invitrogen, A16160) using Immobilon Forte western HRP substrate (Merck, WBLUF0100) and a Chemidoc MP Imaging system (BioRad).

### Hi-C

Hi-C was conducted as described in (Dumont *et al*, 2024) with minor modifications. Briefly, ∼1.5×10^9^ haploid cells were fixed with 3% formaldehyde (Sigma-Aldrich, cat. F8775) for 30 minutes at RT at with orbital agitation at 120 rpm. Formaldehyde was quenched with 330 mM glycine for 20 minutes at RT at 120 rpm. Cells were washed twice with cold water at 3,000 g for 10 minutes. Pellets were split in two tubes and frozen at −80° C. The crosslinked pellets were thawed in ice and resuspended in cold H_2_O supplemented with an anti-protease mix (Roche, cat. 11836170001). Approximately 7.5×10^8^ cells were transferred to a Precellys VK05 tube, and lysed for 3×30s at 6,800 rpm. Between 2 and 5.10^7^ cells were processed for Hi-C using the Arima Hi-C+ kit (Arima Genomics, cat. A410079) following manufacturers’ instructions. The Arima Hi-C+ kit employs a dual restriction digestion (DpnII and HinfI) yielding a median fragment length of 108 bp in *S. cerevisiae*.

DNA was fragmented into 300-400 bp fragments using Covaris M220 sonicator. Preparation of the libraries for paired-end sequencing on an Illumina platform was performed using the Thermofisher Collibri ES DNA Library Prep Kit for Illumina Systems with UD indexes (cat. A38606024) following manufacturer’s instructions. The library was amplified in triplicate PCR reactions using oligonucleotides corresponding to the Illumina sequence adaptors (5’-AATGATACGGCGACCACCGAGATCTACAC-3’ and 5’-CAAGCAGAAGACGGCATACGAGAT-3’) and Phusion DNA polymerase (New England Biolabs, cat. M0531) for 11 cycles. PCR products were purified with AMPure XP beads (Beckman-Coulter, cat. A63881) and resuspended in pure H_2_O. The Hi-C library is quantified using the Qubit DNA high sensitivity kit (Thermo Scientific, cat. Q32851) on a Qubit 2 fluorometer (Thermo Scientific, cat. Q32866). Library quality control, paired-end sequencing (2×150 bp) on Illumina NovaSeq6000 or NovaSeq X Plus, and data QC were performed by Novogene UK. The correspondence between Hi-C libraries and figure panels are listed in **Table EV2**.

### D-loop Capture

D-loop Capture has been performed as described in (Reitz *et al*, 2022; Djeghmoum & Piazza, 2025). The rationale is depicted in **Fig. EV8B**. Briefly, cells were collected at 4°C and resuspended in a solution containing 0.1 mg/mL Trioxsalen and crosslinked upon 365 nm UV irradiation on a BIO-LINK irradiator (Vilber-Lourmat, Cat. BLX-365) for 10 minutes. Cells were spheroplasted and lysed in the presence of 4 pM of APO563 (*i.e.* an oligonucleotide enabling restoration of a restriction site on the resected broken molecule, **Table EV3**). The DNA was digested with EcoRI-HF (NEB, cat. R3101L) and ligated in dilute conditions (≃1.8×10^4^ genome/µL) with T4 DNA ligase (NEB, cat. M0202). DNA was extracted with phenol-chloroform following protein digestion with proteinase K. Psoralen inter-strand crosslinks and adducts were reversed in 100 mM KOH at 90°C for 30 minutes. The pH was neutralized upon addition of 66 mM of NaoAc pH 5.2. Approximately 6 × 10^5^ genome equivalent were used per quantitative PCR (qPCR) reaction, performed in duplicate, on a CFX96 Touch Deep Well Real-Time PCR Detection System (Bio-Rad cat. 3600037), using the SsoAdvanced Universal SYBR Green Supermix (Bio-Rad, cat. 1725274), following manufacturer’s instructions. Primers used are listed in **Table EV3.** qPCR analysis were performed as described in (Reitz *et al*, 2022) using Bio-Rad CFX Maestro and Microsoft Excel.

### Data retrieval from SRA and GEO

Raw paired-end reads from Hi-C experiments in PRJNA526833 (Paldi *et al*, 2020), PRJNA680815 (Jeppsson *et al*, 2022), and PRJNA986466 (Jeppsson *et al*, 2024) were retrieved using the nf-core/fetchngs pipeline (nf-core/fetchngs, 2024). ChIP-Exo profiles from (Rossi *et al*, 2021) (GEO GSE147927 series) were obtained from https://www.datacommons.psu.edu/download/eberly/pughlab/yeast-epigenome-project/.

### Hi-C read alignment and generation of contact maps

Paired-end 150 bp-long reads in fastq.gz format were pre-digested with DpnII and HinfI using parasplit in “all-vs-all” mode. Alternatively, reads were digested using the “cutsite” mode of the Hicstuff *pipeline* function (Matthey-Doret *et al*, 2020a). Hicstuff *pipeline* was used to align pairs of reads independently using Bowtie2 and generate contact data in Graal format (Matthey-Doret *et al*, 2020a, 2022) using a modified *S. cerevisiae* R64-2-1 reference genome in which the *HML, MAT, HMR, URA3, LYS2*, and the right rDNA repeat loci were replaced by “N” of the same length (S288c_DSB_chr3_rDNA reference). In instances where a second *RE* fragment was introduced on chr. IV, the *RE* sequence on chr. III was also masked (S288c_DSB_chr3_rDNA_RE_N reference). Each uniquely mapped read was assigned to a restriction fragment, the uncut, circularization and spurious ligation events were filtered out as described in ref. (Cournac *et al*, 2012) and PCR duplicates removed. The argument -d was set to compute the per-chromosome probability of contact P_c_ as a function of the genomic distance *s* from pairs files. The resulting sparse matrix in Graal format was binned at 1 kb resolution using the Hicstuff *rebin* function and converted to a cooler format with the Hicstuff *convert* function. ICE normalization of the cooler file was performed using the Cooler *balance* function (Abdennur & Mirny, 2020). Iterative coarsening was achieved using the Cooler *zoomify* function, which resulted in a mcool file. Graal and cooler files were used for downstream analysis.

### Generation of Hi-C maps

Hi-C maps were generated from sparse Hi-C data in Graal or cooler format using Hicstuff *view* function, balanced using the SCN method (Cournac *et al*, 2012), log-transformed and binned. Alternatively, ratio maps were generated from cooler files using Serpentine (Baudry *et al*, 2020). Briefly, two matrices of the region of interest binned at 1 kb were subsampled to contain the same number of contacts and converted to a dense format. Comparison of contact maps was performed using the default threshold parameters (50 and 5) following 30 serpentine binning iterations. The trend was set to 0.

### Generation of aggregated Hi-C contact maps

Intra-chromosomal centromere and *GAL* genes pile-ups were generated with Chromosight *quantify* with default parameters (Matthey-Doret *et al*, 2020b) from ICE-normalized Hi-C matrices in cooler format binned at 1 kb and subsampled at 24 million contacts. The pile-ups show the contact enrichment at the coordinates of interests over random genomic sites.

### Computation of the contact probability as a function of genomic distance

Computation of the contact probability as a function of genomic distance P_c_(*s*) and its derivative have been determined from the per-chromosome file generated by Hicstuff *pipeline*. The contact decay probability of the mitochondrial genome and the endogenous 2μ plasmid were removed and the genome average P_c_(s) and slope was computed using the *distance law* function of the Hicstuff package with default parameters within a reference window of 3 and 300 kb.

### Quantification of coverage from Hi-C data

Quantification of coverage from *bam* alignment files generated with the Hicstuff *pipeline* function, which uses Bowtie2 in unpaired mate mode, were sorted and merged with Samtools *sort* and *merged* function, respectively. The coverage was computed in non-overlapping 500 bp bins using Tinycov *covplot* function. The coverage was normalized onto the median genome coverage, and divided over that of samples lacking a DSB on chr. III or chr. IV.

### Quantification of coverage from ChIP-Exo data

Quantification of coverage from *bam* alignment files following sorting with samtools *sort* function were computed as non-overlapping 20 bp bins using Tinycov *covplot* function. The resulting bedgraph files were rendered as histograms or heatmaps on IGV.

### Extraction of intra-chromosomal 4C-like contact profiles

Sparse Hi-C matrices binned at 1 kb in cooler format were loaded, SCN-normalized and converted to dense format using the *flexible_loader* and *sparse_to_dense* function of the Hicstuff package, respectively. Vectors corresponding to the intra-chromosomal contacts made by the 3 kb *RE*-containing region to the right end of chr. III, the 3 kb region flanking the rDNA on the centromere-proximal side toward the left end of chr. XII, and/or the 3 kb region surrounding the insertion point for the *RE* on chr. IV:680,258 were extracted, smoothed with the Savitzky-Golay filter using the *signal.savgol* function of Scipy, and the sum of contacts set to 1. The smoothing kernel size was set to 39 kb with a polynomial order of 2 for whole chromosomes, and to 9 kb with a polynomial order of 1 for zoom on specific features, except otherwise stated. For each viewpoint, 6 control viewpoints located at the same distance from a centromere and on chromosome arms of a similar length were selected, processed in the same way, and their contact frequency averaged. Control viewpoints for the *RE,* in the form chr:coordinate(strand), were V:66000(W), VIII:21000(W), XIV:713000(C), II:153000(W), XI:525631(C), XIII:182646(W). Control viewpoints for the left rDNA-flanking region were IV:747000(C), XIV:332000(W), XIII:565000(C), XV:624000(C), XI:143000(W), II:536000(C). Data (mean ± SEM) of traces corresponding to independent biological replicates are plotted using Graphpad Prism as a distance to the viewpoint. Ratio between two conditions were manually computed under Microsoft Excel.

Computation of the 4C-like contact profiles with the left side of *MAT* as a viewpoint (III:190000) were computed on Hi-C matrices binned at 5 kb. No smoothing was applied.

### Statistical analysis

Statistical tests and linear regressions were performed under Graphpad Prism 10. The experiments were not randomized. The investigators were not blinded to allocation during experiments and outcome assessment. The number of times an experiment has been repeated (n), the nature of the data representation (mean, SEM, etc.), and the statistical test used are indicated in the figure panels or legends. A Student *t* test was used to compare DLC data. Donor preference computed the ratio of D-loops formed at each donor within a single sample. These distributions were compared using an unpaired two-tailed Student *t* test. The absolute DLC data were compared using a paired ratio Student *t* test. This test was chosen because of batch-level variations in D-loop retrieval, which originates from subtle day-to-day differences in crosslinking efficiency (Reitz *et al*, 2022).

### Loop extrusion model

We modeled extrusion as a unidimensional process working along the chromatin: (1) condensin can bind and unbind at a single loading site at rates *k*_*b*_ and *k*_*u*_ respectively; (2) upon binding, one leg of the condensin remains anchored at the binding site while the other translocates along the chromosome at a speed *v*_*e*_. In the idealized scenario of continuous processing without boundaries or roadblocks, the distribution of loop sizes extruded by condensin, *l*, follows an exponential distribution (Brackley *et al*, 2017; Abdulla *et al*, 2022):

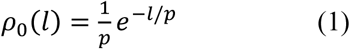

where *p* = *v*_*e*_/*k*_*u*_ is the processivity and represents the average loop size extruded by condensin when it is bound to chromatin. When an impermeable roadblock is present at site *i*, located a genomic distance *l*_*i*_ from the loading site, the loop size distribution is modified as follows:

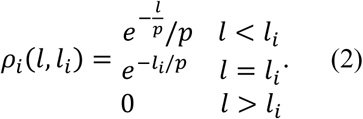

Assuming a density *⍺*_*i*_ of roadblocks at block site *i*, the total loop size distribution for *n* discrete roadblocks is given by

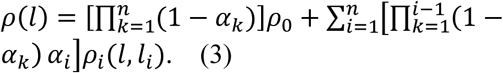

If roadblock sites are not discrete but rather domains of width {Δ*l*_*ki*_}, we assume that one blocking barrier would be positioned randomly within the domain. Therefore, there are 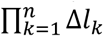 different possible configurations of block site positions. To each possible configuration {*l*_*i*_} corresponds a size distribution (via Eq. (3)). The total loop size distribution is thus the mean value of *ρ*(*l*) averaged over all possible block site configurations.

Given a loop size distribution, we can compute the 4C contact profile between the loading site and other genomic positions using a Gaussian polymer approximation. The contact frequency in the absence of loop extruders is described by (Halverson *et al*, 2014):

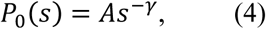

where *A* is a constant, *s* is the genomic distance with the loading site, and *γ* is 1.5 for an ideal polymer and less for more compact regions (Mirny, 2011). In the presence of a fixed loop of size *l*, the contact probability is (Polovnikov *et al*, 2023; Polovnikov & Slavov, 2023):

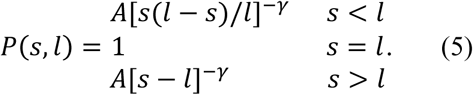

For simplicity we assume *A* = 1. The total 4C profile is then given by:

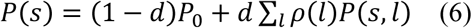

with *d* = *k*_*b*_ /(*k*_*b*_ + *k*_*u*_) the probability to have a condensin bound at the loading site and extruding a chromatin loop.

By adjusting the parameters of the model (*p*, *d*, {*⍺_i_*}, {*l_i_*}, {Δ*l_i_*}), we aim to achieve the best fit with experimental observations. In summary, we compare the ratio of experimental contact frequency to its corresponding control with the theoretical prediction of *P*(*s*)/*P*_0_(*s*). By iteratively adjusting the model parameters, we identify the optimal values. Additionally, we estimate the parameters associated with potential roadblocks by analyzing the heights, lengths, and positions of significant peaks located near these roadblocks. A jupyter python notebook describing the model and allowing to make predictions is available at https://github.com/physical-biology-of-chromatin/CondensinYeast.

## Data availability

The raw Illumina paired-end sequencing and HiC data in cooler format will be made publicly available upon publication.

Software availability:

- Hicstuff (Matthey-Doret *et al*, 2020a) (version 3.2.4 available at https://github.com/baudrly/hicstuff).
- Chromosight (Matthey-Doret *et al*, 2020b) (version 1.6.3, available at https://github.com/koszullab/chromosight).
- Serpentine (Baudry *et al*, 2020) (version 0.1.3, available at https://github.com/koszullab/serpentine).
- Tinycov (version 0.3.1, available at https://github.com/cmdoret/tinycov).
- Integrative Genomics Viewer (Robinson *et al*, 2011) (version 2.8.3, available at https://igv.org).
- Bowtie2 (Langmead & Salzberg, 2012) (version 2.3.5.1 available online at http://bowtie-bio.sourceforge.net/bowtie2/).
- Samtools (Danecek *et al*, 2021) (version 1.3.1 available online at https://github.com/samtools/samtools).
- Cooler (Abdennur & Mirny, 2020) (version 0.9.1 available online at https://github.com/open2c/cooler).
- Parasplit (version 1.1.5 available at https://gitbio.ens-lyon.fr/LBMC/hub/parasplit/).
- Flowing Software (version 2.5.1 freely available online at https://flowingsoftware.com/download/).
- Graphpad Prism (version 10 commercially available at https://www.graphpad.com/).

## Acknowledgements

We thank members of the Piazza, Bernard and Jost laboratories, as well as Armelle Lengronne, Frédéric Beckouët, and Romain Koszul for helpful discussions. We are grateful to Pascal Bernard, Armelle Lengronne, Jim Haber and Stéphane Marcand for their critical reading of the manuscript, and Rodney Rothstein, Thomas Guérin and Stéphane Marcand for sharing yeast strains.

This research was supported by the European Research Council (ERC) under the European Union’s Horizon 2020 (ERC grant agreement 851006) and the Fondation ARC pour la recherche sur le cancer www.fondation-arc.org (ARCPGA2022110005583_6379) to AP, and the Agence Nationale de la Recherche to AP and DJ (ANR-23-CE12-0014-02).

## Author contributions

Conceptualization: AP; Experiments: VP, CD, JS, AD, AP; Data analysis: AP; Modeling: HS, DJ; Data interpretation: HS, DJ, AP; Supervision: DJ, AP; Funding Acquisition: DJ, AP; Manuscript - Writing: AP; Manuscript – Editing: HS, DJ.

## Disclosure and Competing Interests Statement

None.

## Notes

### Competing Interest Statement

The authors have declared no competing interest.

### Summary of Updates

Addition of biological replicates Addition of new HiC data (eg. RE inversion on chr. III and IV) Addition of D-loop capture data showing a functional impact of the condensin-mediated looping in donor selection during recombination. Expanded results and discussion sections

