## Supplementary figures for "Condensin loop extrusion properties, roadblocks, and role in homology search in *S. cerevisiae*"

### Supplemental information

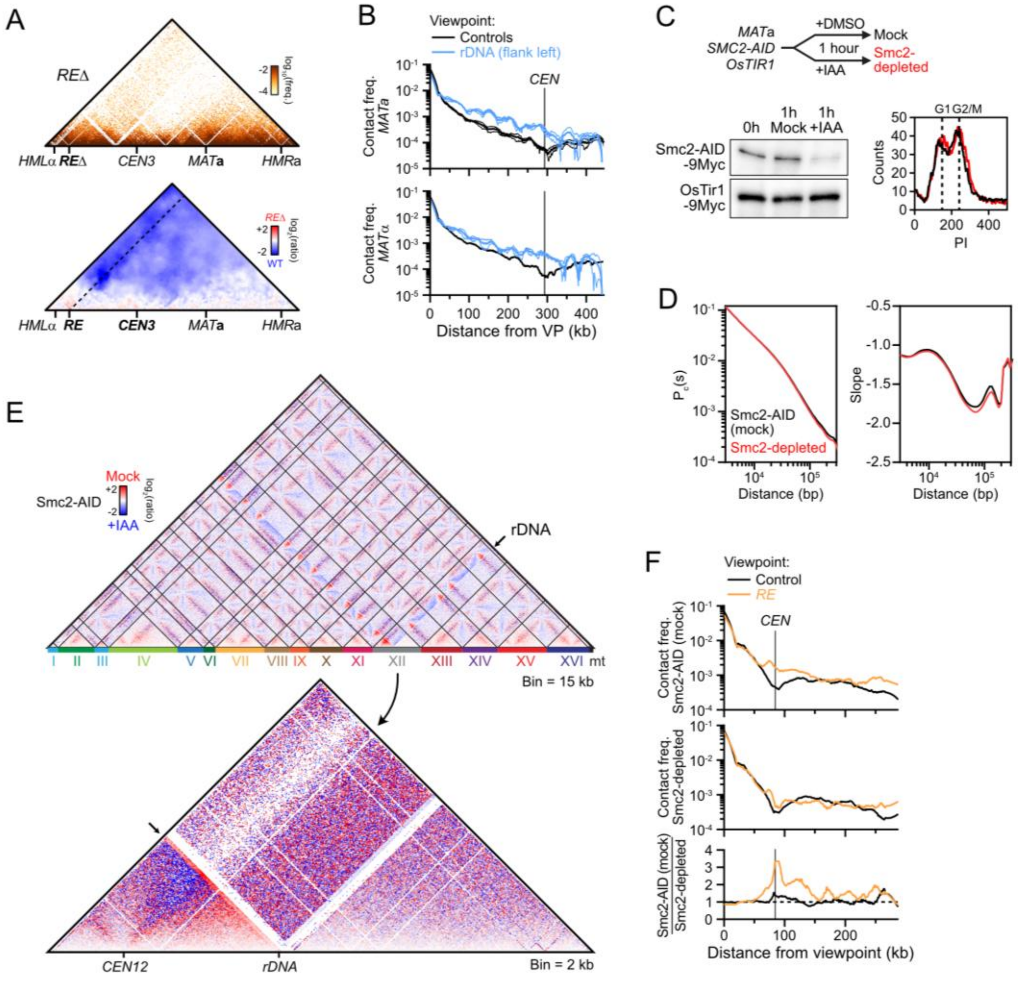


**Figure EV1: Condensin- and *RE*-dependent contact stripes in *MAT*a cells. (related to Figure 1)**

(A) Top: Hi-C contact map of chr. III in a *RE*-deleted *MAT*a strain (APY1548). Bin: 1 kb. Bottom: Ratio map over a WT strain. Data show n=1 biological replicate.

(B) 4C-like contact profile of the left *rDNA*-flanking region (blue) and of the average of 6 control sites (black) as viewpoints in *MAT*a and *MAT*α cells, from Hi-C data in **Fig. 1A**.

(C) Smc2 depletion scheme, Western blot validation, and FACS profiles of Smc2-AID (mock) and Smc2-depleted cells.

(D) Probability of contact as a function of the genomic distance (Pc(s)) and its derivative in Smc2-AID (mock) and Smc2-depleted cells.

(E) Ratio maps of the whole genome (top) and chr. XII (bottom) in cells proficient and deficient for condensin.

(F) Top: 4C-like contact profiles of the *RE* and of the average of 6 control sites in Smc2-AID-tagged (mock) and Smc2-depleted samples, from Hi-C data in **Fig. 1C**. Bottom: Ratio of *RE* and control 4C-like profiles of Smc2-AID (mock) over Smc2-depleted samples. Data show n=1 biological replicate.

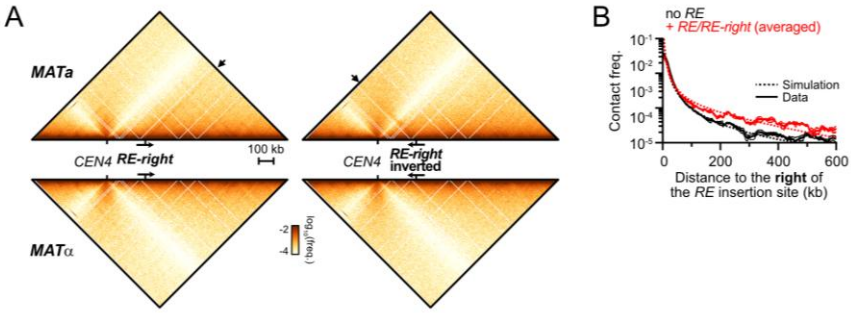


**Figure EV2: Condensin loop extrusion properties. (related to Figure 2)**

(A) Top: Hi-C contact maps of chr. IV in *MAT*a and *MAT*α cells bearing the *RE-right* construct at position 680 kb either in the forward (APY1850 and APY1852) or inverted (APY2058 and APY2060) orientation. Hi-C maps are binned at 5 kb. Data show n=1 biological replicate.

(B) Observed and simulated 4C-like profiles using chr. IV 680 kb as a viewpoint, either unmodified ("no *RE*" black), or upon insertion of the *RE* or the *RE-right* constructs (data averaged). From data in **Fig. 2C**. The ratio of the "+*RE*" profiles over the "no *RE*" profiles gives rise to the normalized data presented in **Fig. 2G**.

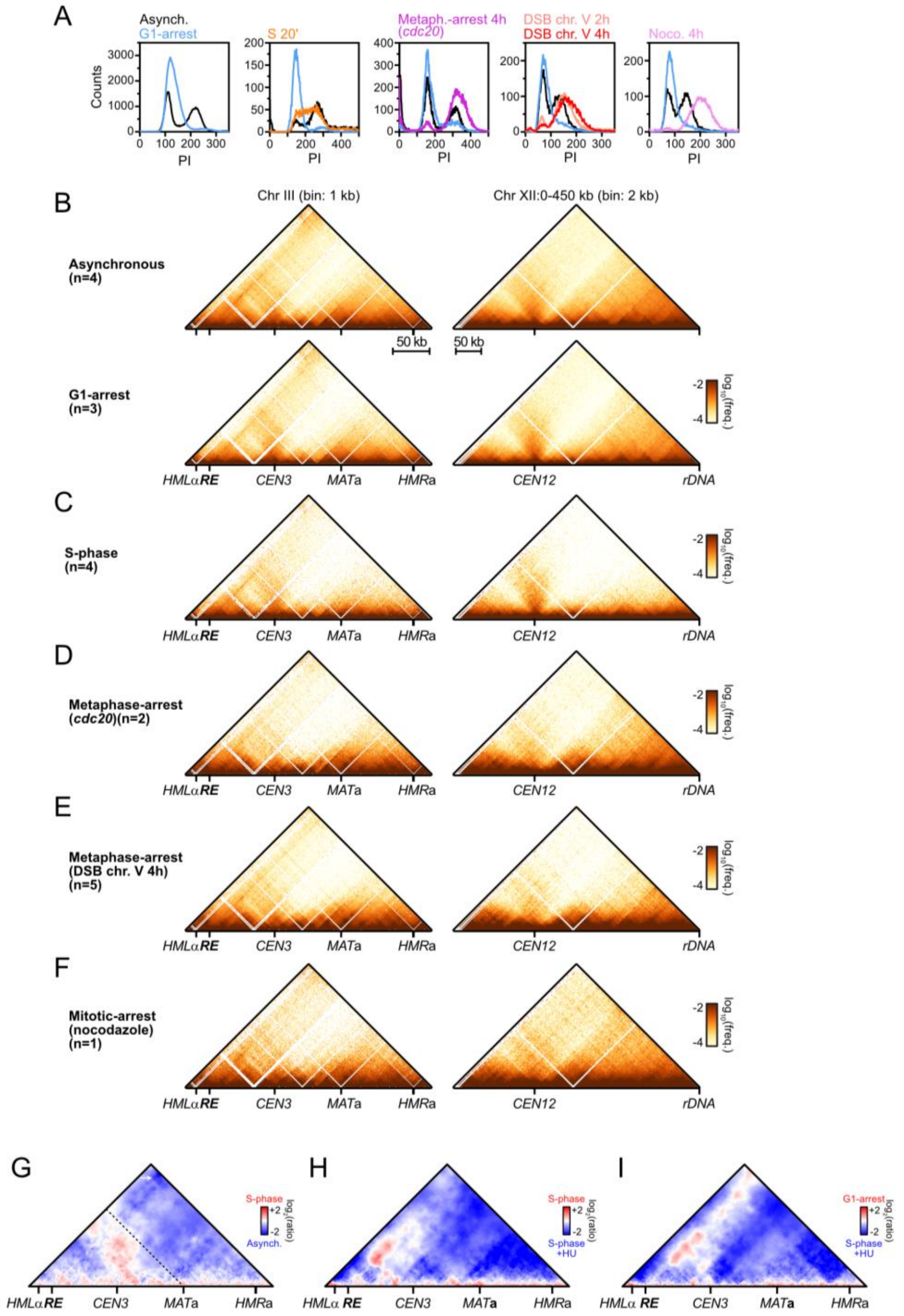


**Figure EV3: Regulation of loop extrusion by condensin across the cell cycle. (related to Figure 3)**

(A) FACS profiles of the different cell-cycle stages studied here.

(B-F) Hi-C maps of chr. III and chr. XII:0-450 kb (A) upon G1-arrest (APY266), (B) during S-phase (APY539 and APY607, merged), (C) upon metaphase-arrest due to *CDC20* repression (APY537), (D) upon DDC-induced metaphase-arrest due to formation of a single unrepairable HO-induced DSB at on chr. V (APY266), and (E) upon mitotic-arrest in the presence of nocodazole (APY266). All cells are

*MAT*a. The number of biological replicates (n) is indicated in each panel. Bin: 1 kb (chr. III) or 2 kb (chr. XII)

(G) Ratio map highlighting the changes to chr. III structure in S-phase vs. asynchronous *MAT*a cells. From data in (B-C).

(H) Same as (G) in untreated vs. HU-treated cells in S-phase. From data in (Jeppsson *et al*, 2022).

(I) Same as (G) in G1-arrested vs HU-treated S-phase cells. From data in (Jeppsson *et al*, 2022).

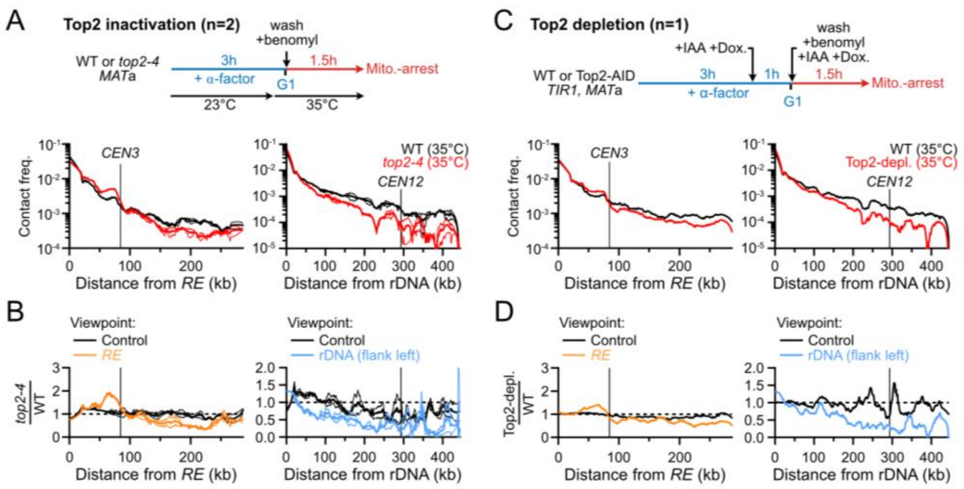


#### Figure EV4: Loop extrusion by condensin is compromised in Top2-deficient cells.

(A) 4C-like profiles of the *RE* (left) and the rDNA (right) and their cognate control sites in a mitotic WT and *top2-4* strains after 1.5 hours at restrictive temperature. Data show mean ± SEM of n=2 biological replicates.

(B) Ratio of 4C-like profiles for the *RE*, *rDNA* and their cognate controls sites in *top2-4* over WT cells. Data show mean ± SEM of n=2 biological replicates.

(C) 4C-like profiles of the *RE* (left) and the rDNA (right) and their cognate control sites in mitotic WT and Top2-depleted strains. Data show n=1 biological replicate.

(D) Ratio of 4C-like profiles for the *RE*, *rDNA* and their cognate controls sites in WT and Top2-depleted cells. Data show n=1 biological replicate.

(A-D) All data are from (Jeppsson *et al*, 2024).

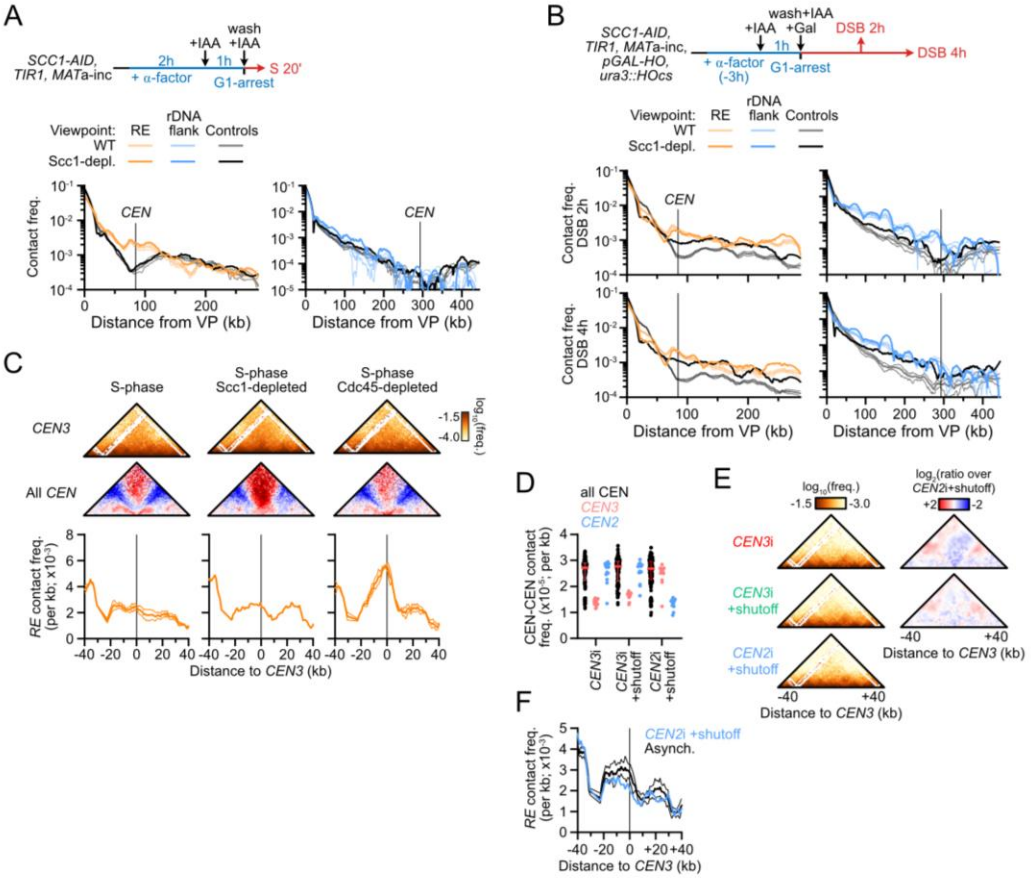


**Figure EV5: The centromere stalls condensin translocation in a kinetochore-dependent manner. (related to Figure 4)**

(A) Loss of Scc1 does not rescue condensin-mediated loop extrusion in S-phase. Top: Scheme for Scc1-AID depletion prior to S-phase release. Bottom: 4C-like profiles at the *RE* and the left *rDNA*-flanking region and their corresponding control sites in WT and Scc1-depleted cells. n = 2 and 1 biological replicates, respectively. Data are from (D'Asaro *et al*, 2025).

(B) Loss of Scc1 does not affect condensin-mediated loop extrusion in metaphase cells. Same as (A), but with Hi-C performed in cells arrested in metaphase upon formation of an unrepairable DSB on chr. V. Note the elevated baseline, particularly at 4 hours post-DSB induction. n = 4 and 1 biological replicates for WT and Scc1-depleted cells, respectively. Data are from (Dumont *et al*, 2024).

(C) Condensin roadblock at *CEN3* in S-phase in WT, Scc1-depleted and Cdc45-depleted cells. Top: Hi-C maps of the *CEN3*-surrounding region. Middle: Aggregated contact maps of all centromeres. Bottom: *RE*-contact stripes. n = 2, 1, and 1 biological replicates for WT, Scc1-depleted and Cdc45-depleted cells, respectively. Data are from (D'Asaro *et al*, 2025).

(D) Inter-chromosomal contact frequency between all centromeres (black), between *CEN3* and other centromeres (pink) and between *CEN2* and other centromeres (blue) following transcription-mediated *CEN3* or *CEN2* inactivation. Each point represents a CEN-CEN contact frequency. Bars show median ± inter-quartile range.

(E) Left: Hi-C contact maps of the *CEN3*-surrounding region (bin: 1 kb). Right: ratio maps of the *CEN3*-surrounding region in *CEN3*-inactivated cells over control *CEN2*-inactivated cells.

(F) 4C-like profiles with the *RE* as a viewpoint in WT and *CEN2*-inactivated strains. Data show mean ± SEM of n=4 and 1 biological replicates, respectively.

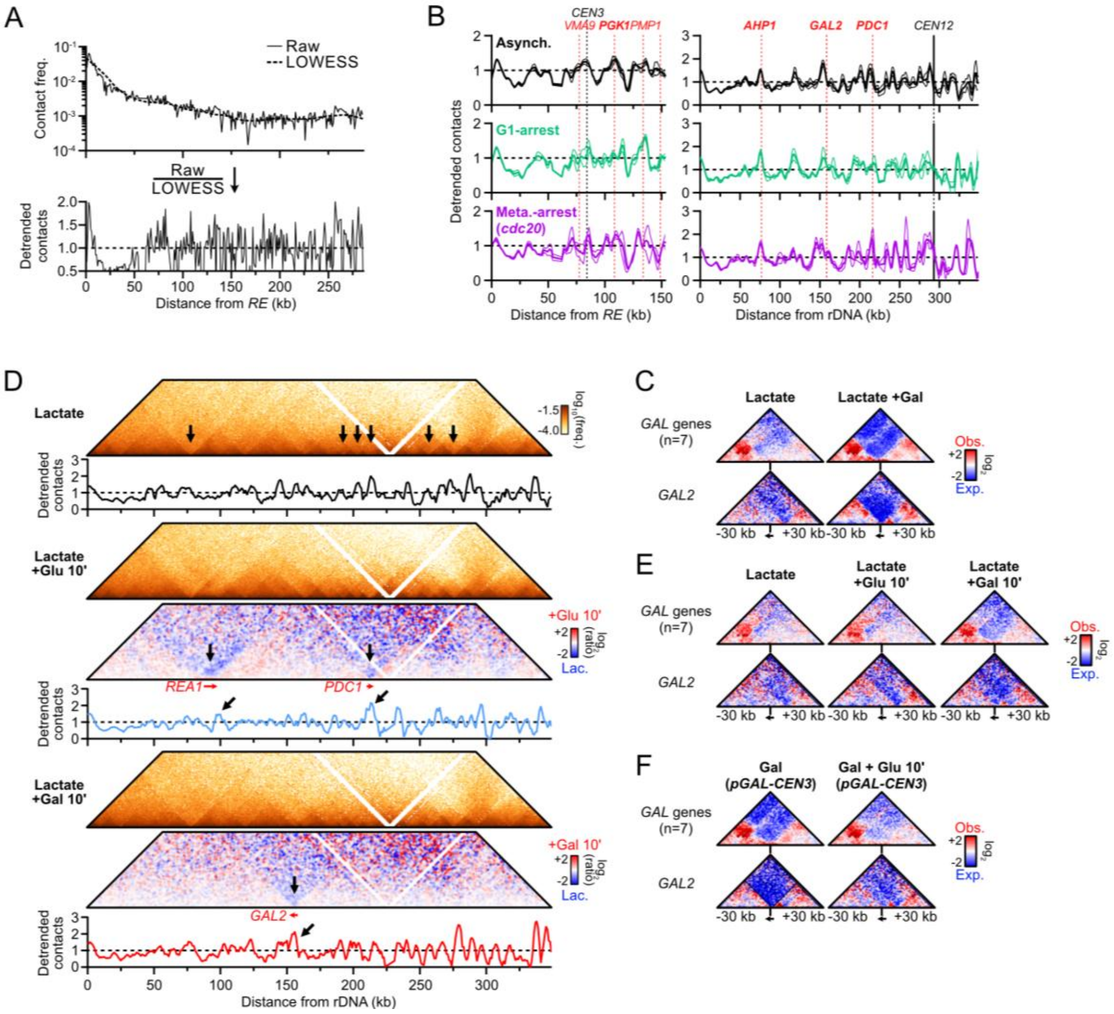


**Figure EV6: Highly transcribed RNA PolII-dependent genes stall condensin translocation. (related to Figure 5)**

(A) Rationale for raw contact detrending over the LOWESS regression.

(B) Detrended *RE* and *rDNA*-flanking contacts in asynchronous, G1-arrested, and metaphase-arrested cells grown in the presence of galactose and in the absence of glucose. Data are the same as in **Fig. 1A** and **3A, C**. Highly-transcribed genes present at the major peaks are indicated.

(C) Observed over expected ratio maps aggregated at all *GAL* genes (top) and at *GAL2* (bottom) in galactose- and glucose-containing media.

(D) Correspondence between high transcription (visible as discrete borders in the Hi-C map) and condensin loop extrusion pausing in lactate media and upon glucose or galactose addition for 10 minutes. n = 1 biological replicate each.

(E) As in (C), from data in (D).

(F) As in (C), upon glucose addition in galactose-containing media. From data in **Fig. 4E** and **5D**.

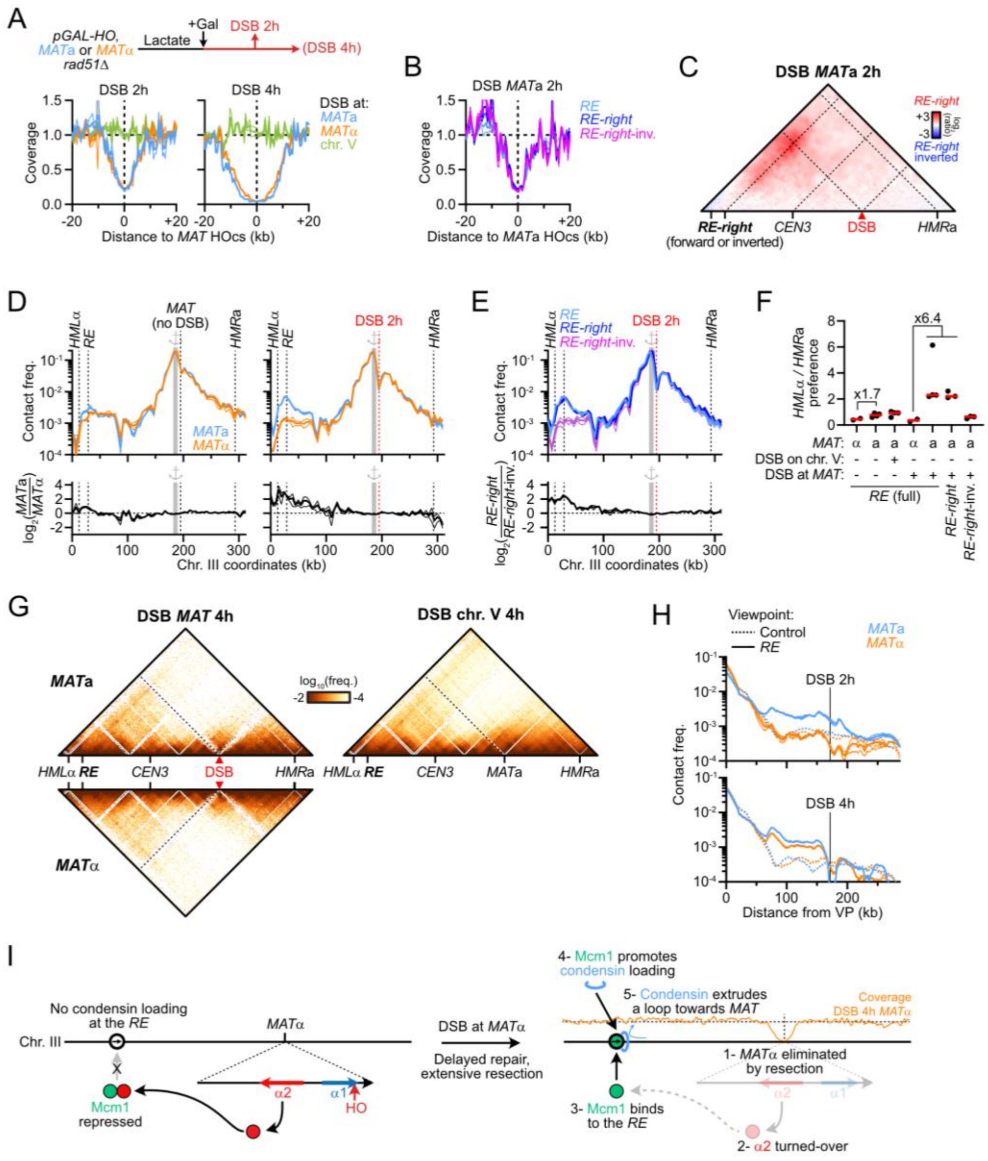

**Figure EV7: DSB formation at *MAT*a blocks condensin translocation and creates a *RE*-DSB loop. (related to Figure 6)**

(A) Coverage from Hi-C reads at 2 and 4 hours post-DSB induction at *MAT* in *MAT*a and *MAT*α cells. Control cells with a DSB on chr. V show no loss of coverage at *MAT*. From data in **Fig. 6A**.

(B) Coverage from Hi-C reads at 2 hours post-DSB induction at *MAT*a in cells with *RE* variants. From data in **Fig. 6D**.

(C) Ratio map of cells with the *RE-right* over the *RE-right*-inverted construct 2 hours post-DSB induction at *MAT*a. From data in **Fig. 6D**.

(D) Top: 4C-like profiles with the 10 kb region left of *MAT* as a viewpoint in asynchronous cells and in cells 2 hours post-DSB induction. Bottom: $Log_2$ ratio of profiles in *MAT*a over *MAT*α cells showing specific enrichment of contact between *MAT*a and the *HML*α-*RE* interval. From data in **Fig. 1A** and **6A**.

(E) Top: 4C-like profiles with the 10 kb region left of *MAT* as a viewpoint in *MAT*a cells bearing different *RE* variants 2 hours post-DSB induction. Bottom: $Log_2$ ratio of 4C profiles in *RE-right* over *RE-right*-inverted-containing cells. From data in **Fig. 6D**.

(F) Quantification of the preference for *MAT* interaction with *HML* vs. *HMR*. Data show individual biological replicates and the median.

(G) Hi-C contact maps in *MAT*a and *MAT*α *rad51*Δ cells (APY1267 and APY1264, respectively) at 4 hours post-DSB induction at *MAT*. Data show n=1 biological replicate each. A *MAT*a cells with an unrepairable DSB on chr. V (APY266) is shown for comparison 2 hours post-DSB induction (n=5 biological replicates). Bin: 1 kb.

(H) 4C-like profiles with the *RE* (or 6 control sites) as a viewpoint, from data in (G) and **Fig. 6A**. Data show mean ± SEM.

(I) Model for the re-activation of condensin-mediated loop extrusion from the *RE* and establishment of a *RE*-DSB loop upon defective repair of a DSB at *MAT*α.

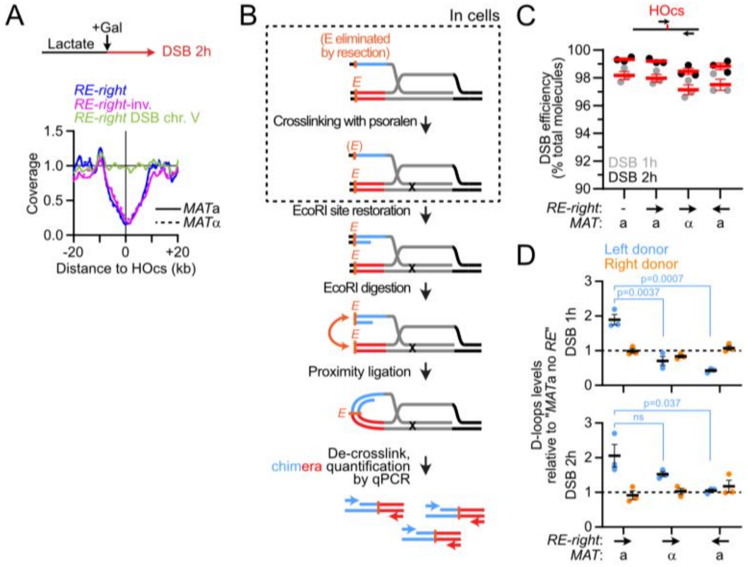


**Figure EV8: The *RE*-DSB loop is portable and promotes *RE*-proximal homology search. (related to Figure 7)**

(A) Coverage from Hi-C reads at 2 hours post-DSB induction at the HOcs on chr. IV in *MAT*a and *MAT*α cells bearing the *RE-right* in forward or inverted orientation. Control *MAT*a cells with a DSB on chr. V show no loss of coverage at that site. From data in **Fig. 7A**.

(B) Rationale of the D-loop Capture assay.

(C) Quantification of DSB formation at 1 and 2 hours post-induction.

(D) D-loops levels expressed relatively to that measured in the *MAT*a strain without *RE* assayed in parallel. From data in **Fig. 7D**. Data show individual biological replicates (n) as well as mean ± SEM. P-values were obtained using a Student *t* test. None of the comparisons for the right donor are significant.
